# Syndecan-4 exerts canonical heparan sulfate-dependent and noncanonical heparan sulfate-independent functions that regulate Aβ amyloid homeostasis

**DOI:** 10.64898/2026.08.21.743990

**Authors:** Kyu Hwan Shim, Yong Ran, Danny Ryu, Brenda D. Moore, Yeeun Yook, Pooneh Amin, Xuefei Liu, Farhana Afroz, Caroline Martin, Mihir Beheray, Wangchen Tsering, Lei Liu, Madison E Platt, Blaine Russell Roberts, Nicholas T. Seyfried, Stefan Prokop, Yona Levites, Todd E. Golde

**Author notes:** Correspondence to Todd E Golde.

## Abstract

**Background:** Heparan sulfate (HS) and heparan sulfate proteoglycans (HSPGs) are components of the amyloid deposits in Alzheimer’s disease (AD) and other amyloidoses. HS and HSPGs are canonically thought to facilitate amyloid deposition by accelerating the aggregation of amyloidogenic proteins and impairing their clearance in a HS-dependent manner.

**Methods:** Leveraging insights from large-scale proteomic data, we focused on Syndecan-4 (Sdc4), the most increased transmembrane HSPG in the AD brain and in the brain of Aβ amyloid depositing mice. We used proximity ligation assays (PLA) to evaluate the association of Sdc4 with Aβ *in situ* and assessed the impacts of the Sdc4 ectodomain on Aβ aggregation *in vitro*. Overexpression studies in cells, hiPSC-derived neurons, and mouse organotypic brain slice cultures (OBSCs) coupled with structure-function studies were used to investigate impacts on Aβ production and APP processing. Finally, effects of overexpression of Sdc4 *in vivo* in the CRND8 amyloid deposition model were evaluated.

**Results:** Consistent with canonical roles, PLA demonstrated a spatial association of Sdc4 with amyloid deposits, and *in vitro*, the Sdc4 ectodomain accelerated Aβ fibril formation in a HS-dependent manner. Unexpectedly, Sdc4 overexpression reduced Aβ production in CHO cells, hiPSC-derived neurons, and OBSCs. These effects were accompanied by dramatic decreases in the levels of sAPPα and C83 and increased immature APP in the cell. Sdc4 promoted altered APP localization into detergent resistant membrane domains and increased APP association with ATG5+/LC3+/Cathepsin D+ vesicles. Structure-function studies revealed that the transmembrane region mediates these effects in a glycosaminoglycan-independent manner. Sdc4 overexpression in the brain of APP mice significantly reduced amyloid deposition at an early age.

**Conclusions:** Sdc4 exerts paradoxical and mechanistically distinct effects that could impact AD pathogenesis differentially, potentially promoting Aβ fibrillization extracellularly while suppressing APP processing and Aβ production. Such data challenge the prevailing view that increased levels of HSPGs in AD are always pro-amyloidogenic and identify Sdc4 as a previously unrecognized regulator of amyloid homeostasis in AD.

**Graphical Abstract:** 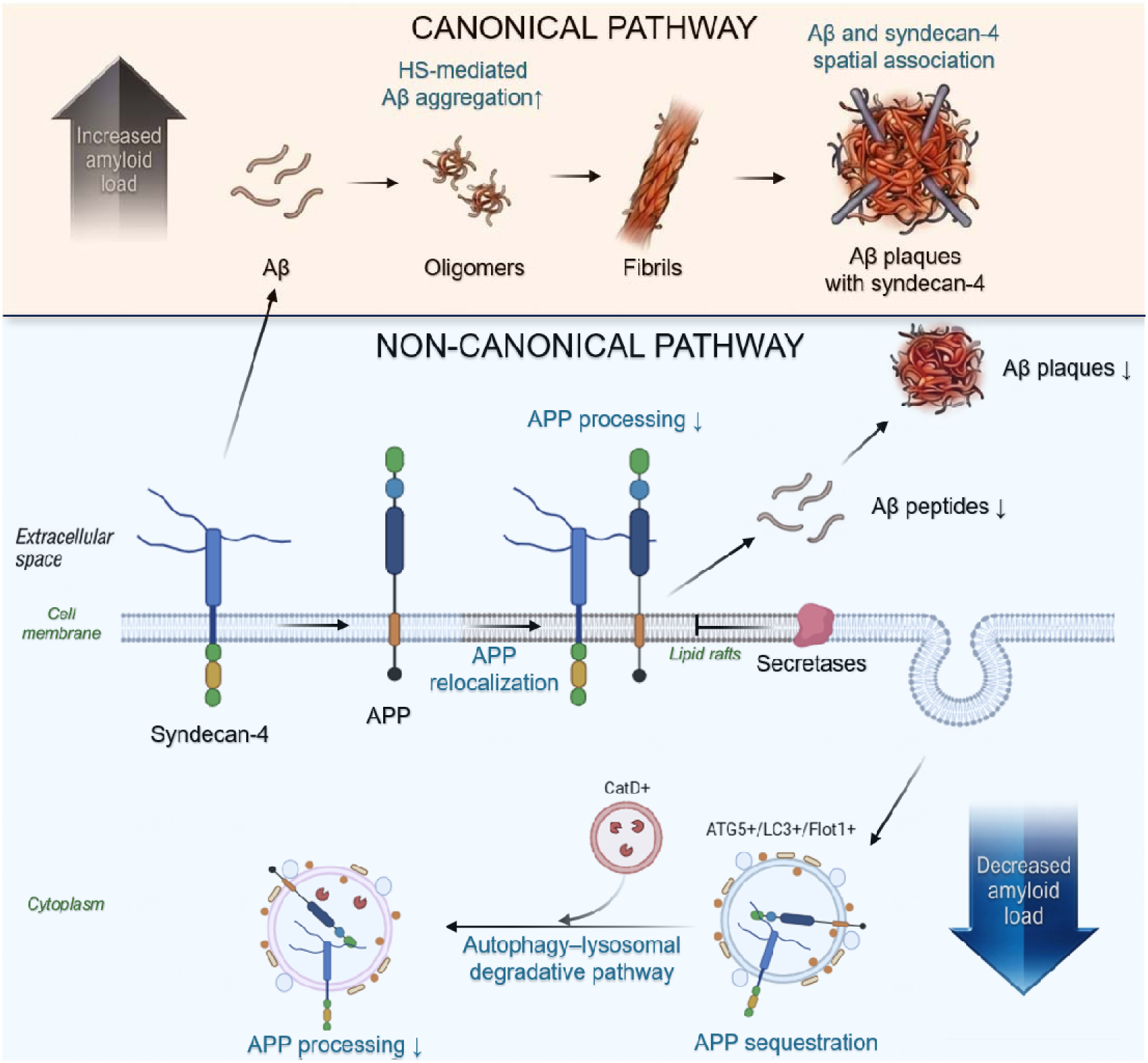

## Background

Alzheimer’s disease (AD) is a progressive neurodegenerative disorder characterized by gradual cognitive decline, synaptic dysfunction, and widespread neuronal loss [1–3]. Key pathological features include the accumulation of amyloid-β (Aβ) and the formation of neurofibrillary tangles (NFTs) composed of hyperphosphorylated tau protein. Extensive research has focused on Aβ pathology, leading to the development of therapeutic strategies targeting Aβ production, aggregation, and clearance [4, 5]. Aβ plaques are embedded within, and appear to disrupt, the extracellular matrix (ECM) in AD. Indeed, numerous ECM proteins are among the most increased proteins in the AD brain and in mouse models of amyloid deposition [6–14]. Many of these proteins are heparan sulfate proteoglycans (HSPGs) themselves or are known to bind HS or HSPGs. Central to this milieu are HSPGs, long recognized as important constituents of Aβ amyloid plaques and other amyloid deposits [15]. In the context of AD, agrin (AGRN) and perlecan (HSPG2) have been the most intensively studied HSPGs [16, 17], though more recent work has explored HSPGs more broadly, including glypicans (GPCs), SPARC (osteonectin), CWCV and kazal-like domain proteoglycans (SPOCK1-3), and Syndecans (SDC1-4) [18–29].

Neuropathological studies have shown that multiple HSPG family members, including agrin, syndecans, and glypicans, are enriched within amyloid plaques and co-localize with fibrillar Aβ deposits [30–32]. In addition, heparan sulfate (HS) has been shown to directly promote Aβ fibrillization and accelerate its aggregation kinetics [33]. HSPGs have also been implicated in NFT pathology, as accelerants of tau fibrillization and as possible receptors for tau internalization [20, 34–39]. Thus, HSPGs that accompany amyloid deposition represent a possible link between amyloid and tau pathology. Beyond AD, HSPGs have also been implicated in other amyloid-related disorders, where they are again proposed to interact with amyloidogenic proteins, promote fibril formation, and contribute to the stabilization and persistence of amyloid deposits [40].

Proteomic studies of human AD brains have increasingly characterized the molecular composition of the amyloid plaque microenvironment, identifying a diverse set of proteins enriched in and around Aβ plaques and establishing that plaques have highly organized molecular environments [6, 41]. Analysis of these datasets has revealed a significant enrichment of extracellular matrix-associated proteins, including components of the matrisome, a conserved protein network closely linked to amyloid pathology [6, 13]. Among these, HSPGs have been consistently implicated in AD pathology [31]. Many proteins that co-accumulate with amyloid deposits have been shown to bind amyloid fibrils; however, a substantial proportion are also known to interact with HS and HSPGs [16]. These observations suggest the existence of complex, multipartite interactions among amyloid fibrils, HS/HSPGs, and other co-accumulating proteins. Although HS has been proposed to facilitate amyloid nucleation and stabilize fibrils, the high prevalence of HS-binding proteins within the amyloid-associated proteome complicates the delineation of causal relationships. Current evidence indicates that both amyloid-mediated and HS-mediated interactions contribute to the co-accumulation of proteins within amyloid deposits.

Syndecans are a family of four transmembrane HSPGs that function as co-receptors for heparin-binding growth factors and play key roles in cell signaling, adhesion, and extracellular matrix interactions [42, 43]. Postmortem analyses of human AD brains have reported increased expression of syndecans and their co-localization with amyloid plaques [6, 32, 44]. Overexpression studies have shown that syndecans facilitate the cellular uptake and fibrillization of Aβ_42_ [45]. Furthermore, syndecans mediate the cellular uptake of Aβ and tau fibrils via a lipid raft-dependent, clathrin-independent endocytic pathway [45, 46]. As we previously reported, syndecan-4 (Sdc4) was a core member of the cross-species conserved M42 matrisome network enriched within amyloid plaques in the brains of AD patients and CRND8-Tg mice overexpressing amyloid precursor protein (APP) with Swedish and Indiana mutations [6]. In addition, Sdc4 serves as a receptor for midkine and pleiotrophin, two heparin-binding growth factors that promote Aβ fibrillization and plaque deposition. Despite these associations, the specific contribution of Sdc4 to Aβ-related pathways remains poorly defined. Given its established roles in membrane trafficking, endocytosis, and cytoskeletal organization, Sdc4 may influence Aβ binding at the cell surface and its subsequent intracellular pathway, yet whether and how Sdc4 actively participates in Aβ pathology in AD remains unresolved.

In this study, we investigated the domain-dependent functions of Sdc4 in amyloid biology. Using complementary *in vitro*, *ex vivo*, and *in vivo* approaches, we examined how the extracellular and intracellular domains of Sdc4 differentially regulate Aβ aggregation and APP processing and assessed the net effect of Sdc4 overexpression on amyloid pathology.

## Methods

### Proteomics data analysis

Previously generated in-house proteomics datasets[6] from 12-and 18-month-old CRND8-Tg mice and AD patient samples were reanalyzed to identify differentially expressed proteins (DEPs). Volcano plots were generated to visualize DEPs within each dataset, and shared DEPs between CRND8-Tg mice and AD patient samples were identified by direct intersection of significant protein lists.

### Construct design

All cDNAs used in this study were synthesized by GenScript (Piscataway, NJ) and subcloned into the recombinant adeno-associated virus based on serotype 2 (rAAV2) vector with CBA promoter (pCTR1) [47]. The APP-GFP construct (pEGFP-N1-APP, #69924) was obtained from Addgene. All constructs, except amyloid precursor protein containing the Swedish mutation of K595N and M596L (APPswe) and GFP, included a C-terminal FLAG tag (DYKDDDDK). The extracellular domain construct of syndecan-4 (eSdc4) comprised amino acids 1–145, lacking the transmembrane and cytoplasmic regions. The Sdc4 C-terminal fragment (Sdc4-CTF) included amino acids 136–198, and the truncated Sdc4-178 construct contained amino acids 1–178. The Sdc4 transmembrane domain (Sdc4-TMD) construct was engineered to encompass the transmembrane domain (amino acids 136–156). A point mutation was introduced within this domain, replacing glycine at position 157 with leucine (G157L). For organotypic brain slice cultures (OBSCs), constructs were re-cloned under the human synapsin1 promoter to achieve neuron-specific expression, and the GFP construct under the same promoter included a C-terminal FLAG tag.

### Production of recombinant eSdc4 in HEK293T cells

HEK293T cells were maintained in Dulbecco’s Modified Eagle Medium (Corning, #17-207-CV) supplemented with 10% FBS (Avantor Seradigm, #76419-584) and 1% penicillin/streptomycin (PS; Thermo Fisher Scientific, #15140122) at 37 °C in a humidified incubator with CO_2_. Cells were seeded in culture dishes and transfected at 70-80% confluency with the eSdc4 expression plasmid using polyethyleneimine. After overnight incubation, the culture medium was replaced with Opti-MEM (Thermo Fisher Scientific, #31985070), and the cells were cultured for an additional two days before collection of the conditioned medium. Conditioned medium containing secreted recombinant eSdc4 was collected after 48 hours of incubation in Opti-MEM. The medium was centrifuged at 3,000 x g for 10 minutes at 4 °C to remove cell debris. The supernatant was concentrated using a 10-kDa molecular weight cutoff ultrafiltration device (MilliporeSigma, #PLGC07610) to approximately 1/20 of the original volume. Anti-DYKDDDDK G1 affinity resin (GenScript, #L00432) was used to capture recombinant eSdc4-FLAG. Then, proteins were eluted using IgG elution buffer (Thermo Fisher Scientific, #21009), and the eluate was immediately neutralized with 1 M Tris buffer (pH 7.5). Purified proteins were buffer exchanged into PBS using spin desalting columns (Thermo Fisher Scientific, #89892). The purified proteins were stored at −80□°C until use.

### Thioflavin T fluorescence assay for Aβ

Aβ_40_ peptide (Anaspec, #AS-24236) was solubilized in 1,1,1,3,3,3-Hexafluoro-2-propanol (HFIP; MilliporeSigma, #52517) and dried overnight in a fume hood. HFIP-treated Aβ_40_ or Aβ_42_ peptide (MilliporeSigma, #AG968) was dissolved in 1% NH_4_OH and subsequently diluted in PBS. Monomeric Aβ (2.5 µM) was incubated with or without eSdc4 in the presence of 50 µM Thioflavin T (ThT) in black 96-well plates. Samples were maintained at 37 °C, and fluorescence intensity was measured every 10 minutes for up to 24 hours using a Molecular Devices FlexStation plate reader, with excitation at 440 nm and emission at 485 nm. Plates were shaken for 15 seconds between each read. A fluorescence threshold was determined as the mean of the lowest 10 baseline values plus ten times their standard deviation. Lag time was defined as the first time point at which the fluorescence signal exceeded this threshold. The data were averaged over every three consecutive time points and plotted using GraphPad Prism.

### Transmission electron microscopy (TEM)

Morphology of Aβ_42_ aggregates was assessed by collecting aliquots at each time point from the ThT kinetic assay for transmission electron microscopy (TEM). Samples were applied to glow-discharged, carbon-coated copper grids and incubated for 1-2 minutes, after which excess liquid was blotted off with filter paper. Grids were then negatively stained with 2% (w/v) uranyl acetate for 45–60 seconds, followed by gentle blotting and air-drying. TEM imaging was performed at 80 kV using a transmission electron microscope.

### Preparation of Aβ monomers and fibrils

Aβ species were initially dissolved in 1% NH₄OH and subsequently diluted in PBS. Monomeric

Aβ_42_ was used immediately for binding analysis. To generate Aβ_42_ fibrils, Aβ_42_ (AnaSpec, #AS-24225) was incubated at 37 °C with gentle shaking overnight. Aβ_40_ was incubated under the same conditions at 37 °C for 1 week. Following incubation, the samples were centrifuged at 14,000 x g for 10 minutes at room temperature to pellet the fibrils. The supernatant was carefully removed, and the fibril pellet was resuspended in PBS and stored at −80□°C until use.

### Aβ immobilization-based binding assay

Plates were coated with monomer or fibrillar forms of Aβ species. Aβ was diluted to 2 μg/mL in PBS following brief probe sonication (2 s, 50% amplitude) to ensure proper dispersion. A total of 100 μL of the diluted Aβ solution was added to each well, and plates were sealed and incubated overnight at 4°C. Prior to sample incubation, wells were incubated with blocking buffer for 2 hours at room temperature. After removal of wash buffer, 100 μL of the samples were added to each well. Plates were sealed and incubated overnight at 4°C. For detection, FLAG-HRP conjugate (MilliporeSigma, #A8592) was added to each well. Plates were incubated for 2 hours at room temperature. Following incubation, TMB substrate solution (Thermo Fisher Scientific, #34028) was added, and the plates were incubated for 30 min at room temperature in the dark. The reaction was stopped by adding 100 μL stop solution to each well. Signal was subsequently measured using the plate reader.

### Cellular transfection and sample collection

CHO cells were maintained in Ham’s F-12 medium (Cytiva, #SH30026.01) supplemented with 10% FBS and 1% PS. H4 cells were cultured in Opti-MEM medium supplemented with 5% FBS and 1% PS. CHO cells were transfected using Lipofectamine LTX Plus reagent (Thermo Fisher Scientific, #15338100), and H4 cells were transfected using FuGENE HD (Promega, #E2311), following the manufacturer’s protocols. After overnight incubation with transfection complexes, the medium was replaced with Opti-MEM, and cells were cultured for an additional 24 hours. The cell supernatant was harvested and centrifuged at 1,000 x g for 10 minutes at 4°C to remove debris, followed by supplementation with protease inhibitor cocktail (Roche, #11697498001). Cell lysates were prepared in cytoskeleton (CSK) lysis buffer composed of 25 mM Tris-HCl (pH 7.5), 150 mM NaCl, 1 mM EDTA, 1% Triton X-100 (TX-100), and 20 mM NaF, with protease inhibitors. Cells were lysed in this buffer and subjected to probe sonication to ensure complete disruption. Lysates were then centrifuged at 16,000 x g for 20 minutes at 4°C, and the supernatants were collected. All samples were stored at −80°C until further analysis.

### Detergent-based APP fractionation

HEK293T cells were seeded in 6-well plates to reach 70% confluency at the time of transfection. Cells were transfected using polyethylenimine with the following conditions: APP_swe_ + empty vector control or APP_swe_ + Sdc4. Cells were lysed in ice-cold CSK buffer containing 1% Triton X-100 (TX-100) and protease inhibitors, then incubated for 30 min at 4°C. Lysates were centrifuged at 16,000 x g for 20 min at 4°C, and the supernatant was collected as the TX-100 soluble fraction (non-raft-associated). The remaining pellet was washed once with ice-cold PBS and centrifuged again. The pellet was resuspended in RIPA buffer containing 1% SDS and protease inhibitors. Samples were then probe-sonicated for 5 seconds at 50% power to ensure complete solubilization, followed by centrifugation at 16,000 x g for 20 min. The supernatant was collected as the detergent-resistant membrane (DRM, raft-associated) fraction. Samples were mixed with loading buffer and heated at 70°C for 5 min before SDS-PAGE. APP was detected by immunoblotting using an anti-APP antibody.

### ELISA

Aβ species in culture medium and brain homogenates were quantified using sandwich ELISA, as described previously [48]. Total Aβ was captured with antibodies mAb5 or mAb9 (Human Aβ_1-16_; T.E. Golde) and detected with HRP-conjugated mAb 4G8 (Human Aβ_17-24_; BioLegend, #800720). To measure Aβ_1-40_, samples were captured with HRP-conjugated mAb 13.1.1, which specifically recognizes human Aβ_35-40_ (T.E. Golde). For Aβ_1-42_, mAb 2.1.3 (Human Aβ_35-42_; T.E. Golde) was used for capture. HRP-conjugated mAb 33.1.1 (Human Aβ_1-16_; T.E. Golde) served as the detection antibody for both Aβ_1-40_ and Aβ_1-42_. Optical densities were recorded using a microplate reader. For brain homogenates, Aβ levels were normalized to the total protein content of the corresponding lysates.

### Aβ Immunoprecipitation and MALDI-TOF Mass Spectrometry

Stable cell supernatant samples were quantified by matrix-assisted laser desorption/ionization time-of-flight mass spectrometry (MALDI-TOF MS) following a modified protocol.[49] Each 500LµL sample was combined with 500 µL of binding buffer (100 mM Tris-HCl, pHL7.4; 300LmM NaCl; 0.2% n-dodecyl-β-D-maltoside; 0.2% n-nonyl-β-D-thiomaltoside) containing 10% N4PE CSF diluent (Quanterix). A total of 12 µg of Dynabeads M-270 Epoxy (Thermo Fisher Scientific, #14301) pre-coupled with anti-Aβ antibodies (6 µg mAb5) was added, and the mixture was incubated at 4 °C for 1.5 hours with rotation. Following immunoprecipitation, the beads underwent sequential washes with binding buffer plus 10% N4PE, binding buffer alone, PBS twice, and HPLC-grade water, with transfers to fresh Protein LoBind tubes to minimize non-specific carryover. Peptides were eluted in 6 µL of 3 mg/mL α-cyano-4-hydroxycinnamic acid (CHCA; Bruker) dissolved in TA50 solvent (50% acetonitrile, 0.1% trifluoroacetic acid, 1LmM ammonium dihydrogen phosphate). One microliter of the eluate was spotted in duplicate on a Bruker AnchorChip target plate and air-dried at room temperature. MALDI-TOF MS spectra were collected on a Bruker NeoFlex instrument in positive reflector mode across an m/z range of 3,500–5,500, accumulating 5,000 laser shots per spot at roughly 40% laser power. Peptide standards were used for external calibration. FlexControl software (Bruker) handled spectral processing, including Savitzky-Golay smoothing and baseline subtraction. Peak areas corresponding to Aβ species (Aβ_37_, Aβ_38_, Aβ_40_, and Aβ_42_) were measured for quantification.

### Western blot analysis

The samples were denatured by heating in loading buffer at 55L°C for 5 minutes. Proteins were separated on 4–12% Criterion™ XT Bis-Tris Protein Gel (Bio-Rad, #3450123) and transferred onto PVDF membranes for 7 minutes using Trans-Blot Turbo Transfer System (Bio-Rad, #1704150). Membranes were blocked for 1 hour at room temperature, followed by overnight incubation at 4L°C with primary antibodies targeting sAPPα (1:2000; mAb5), sAPPβ (1:1000; Biolegend, #813401), C-terminus APP (1:1000; MilliporeSigma, #171610 or #A8717), FLAG (Biolegend, #902401), β-tubulin (1:2000; Bio-Rad, #VMA00453), or β-tubulin III (1:1000; Thermo Fisher Scientific, #14-4510-82). After washing with TBS containing 0.1% Tween-20 (TBST), membranes were incubated with fluorophore-conjugated secondary antibodies (LICORbio, IRDye® 680RD Goat Anti-Mouse IgG or IRDye® 800CW Donkey Anti-Rabbit IgG) for 1Lh at room temperature. Following three additional washes with TBST, the membranes were imaged using an Odyssey Infrared Imaging System (LI-COR Biosciences).

### PLA

Proximity ligation assay (PLA) was performed using the Duolink *in situ* PLA Kit (MilliporeSigma, #DUO92007) according to the manufacturer’s instructions with minor modifications. Cultured cells were washed with PBS and fixed with 4% paraformaldehyde for 15 minutes, followed by permeabilization with 0.1% Triton X-100 in PBS for 10 minutes. To detect only proteins on the cell surface, primary antibodies were incubated for 1 hour at 4 °C, followed by fixation without permeabilization. For paraffin-embedded brain sections, slides were deparaffinized in three changes of xylene (5 minutes each) and rehydrated through a graded ethanol series (100%, 100%, 95%, and 70%). Slides were then washed in distilled water and subjected to antigen retrieval by steaming in distilled water for 30 minutes. Sections were subsequently incubated in PBS 0.1% with Tween-20 for 30 minutes at room temperature and washed with Wash Buffer A. The samples were then blocked with Duolink blocking solution for 1 hour at 37 °C. For the control group, the binding site of mAb5 was blocked by incubating with Aβ_1–16_ peptide for 1 hour at 37 °C. Each pair, consisting of anti-Sdc4 antibody (T.E. Golde) and one of the antibodies targeting Aβ_1–x_ (mAb5), Aβ_1–x_ (82E1), Aβ_x–40_ (13.1.1), or Aβ_x–42_ (2.1.3) was diluted in Duolink antibody diluent and incubated overnight at 4 °C in a humidified chamber. After washing with Wash Buffer A, cells were incubated with PLA probe PLUS and MINUS antibodies diluted 1:5 in antibody diluent for 1 hour at 37 °C. Ligation was then performed using ligation buffer containing ligase for 30 minutes at 37 °C, followed by rolling-circle amplification using amplification buffer containing polymerase for 100 minutes at 37 °C. After amplification, samples were washed with Wash Buffer B and briefly rinsed with diluted Wash Buffer B. Slides were mounted using Duolink mounting medium containing DAPI (MilliporeSigma, #DUO82040) and imaged using a fluorescence or confocal microscope.

### Immunofluorescence

Transfected cells were washed with PBS and fixed with 4% paraformaldehyde for 15 minutes at room temperature. Cells were permeabilized with 0.1% Triton X-100 in PBS for 10 minutes and blocked with 5% bovine serum albumin (Thermo Fisher Scientific, #BP1600-100) in PBS for 1 hour at room temperature. Primary antibodies were diluted in blocking buffer and incubated with the cells overnight at 4 °C. The following antibodies were used: anti-Aβ_1-16_ (mAb5, T.E. Golde), anti-APP (Millipore Sigma, #MAB348), anti-ATG5 (Cell Signaling Technology, #2630), anti-LC3B (Cell Signaling Technology, #3868), anti-cathepsin D (Abcam, #ab6313), anti-flotillin (Cell Signaling Technology, #18634S) and anti-FLAG. After washing with 0.05% Tween-20 in PBS, cells were incubated with species-appropriate fluorophore-conjugated secondary antibodies for 1 hour at room temperature in the dark. Cells were mounted using antifade mounting medium with DAPI (Vector Laboratories, #H-2000-10). Fluorescence images were acquired using a fluorescence microscope.

### AV production

rAAV2/8-hSyn1-APPswe, rAAV2/8-hSyn1-GFP, or rAAV2/8-hSyn1-Sdc4 used for OBSCs were produced and purified as previously described.[50] Since hiPSC-derived neurons are more efficiently transduced by the AAV9-based vector, rAAV2/9P31-hSyn1-GFP and rAAV2/9P31-hSyn1-Sdc4 were generated. AAV vectors containing either the cytomegalovirus enhancer/chicken β-actin (CBA) promoter or the human synapsin 1 promoter, along with a woodchuck hepatitis virus post-transcriptional regulatory element (WPRE) and a bovine growth hormone polyadenylation signal, were produced in HEK293T cells using PEI-mediated transfection. Cells were co-transfected with AAV helper plasmids (DP8.ape). Seventy-two hours post-transfection, cells were collected and lysed in buffer containing 0.5% sodium deoxycholate and 50 U/mL Benzonase (MilliporeSigma, # 71206-3) through multiple freeze-thaw cycles at −80 °C and 50 °C. Viral particles were subsequently purified using a discontinuous iodixanol gradient. Samples were exchanged into PBS using a 100 kDa molecular weight cutoff centrifugal filter unit (MilliporeSigma, #UFC910008). Viral genomic titers were quantified by qPCR using a CFX384 system (Bio-Rad). Viral DNA was prepared by DNase I treatment (Thermo Fisher Scientific, #18068015), followed by heat inactivation, protein digestion with Proteinase K, and a second heat inactivation step. Freshly prepared AAVs were aliquoted and stored at −80 °C.

### Differentiation of hiPSC-derived forebrain neurons

Human induced pluripotent stem cells (hiPSCs) were maintained in mTeSR™ Plus medium (STEMCELL Technologies, #100-1130) on matrigel-coated plates under standard culture conditions. For neural induction, hiPSCs were dissociated into single cells using Accutase (STEMCELL Technologies, #07920) and plated in STEMdiff™ Neural Induction Medium (STEMCELL Technologies, #05839) according to the manufacturer’s instructions. Cells were cultured for 7 days with daily medium changes to generate neural progenitor cells (NPCs). NPCs were expanded through three passages and subsequently differentiated into forebrain neurons using the STEMdiff™ Forebrain Neuron Maturation Kit (STEMCELL Technologies, #08605). On day 1 of maturation, neurons were transduced with either rAAV2/9P31-hSyn1-GFP or rAAV2/9P31-hSyn1-Sdc4, with medium changes every 2-3 days.

### Organotypic brain slice culture

OBSCs were prepared from postnatal day 8-9 B6/C3H mice as previously described.[51] Briefly, pups were cryoanesthetized and decapitated, and the brains were rapidly removed and dissected in sterile ice-cold dissection buffer consisting of HBSS without calcium and magnesium (Thermo Fisher Scientific, #14175079), supplemented with 2 mM ascorbic acid (STEMCELL Technologies, #72132), 39.4 μM ATP (MilliporeSigma, #A6419), and 1% PS. Tissues were then maintained in Basal Medium Eagle supplemented with 0.5 mM ascorbic acid, 26.6 mM HEPES, Glutamax, 0.033% Insulin, 0.5% PS, and 25% horse serum. Coronal brain slices (300 μm thickness) containing cortex and hippocampus were prepared and plated onto semi-porous membrane inserts (MilliporeSigma, 0.4 μm pore diameter) in 24-well culture plates, with one slice per insert. Slices were maintained in slice culture medium under standard culture conditions.

For viral transduction, slices were treated at the start of culture (0 days *in vitro*, DIV) with the indicated rAAVs. Slices were co-transduced with rAAV2/8-hSyn1-APPswe together with rAAV2/8-hSyn1-Sdc4 or their respective GFP control vectors. Viral particles were applied directly to the culture medium at a total dose of approximately 1 x 10^14^ genome particles per well. Transduction efficiency was verified by GFP fluorescence imaging prior to sample collection. After 10-14 days in culture, slices were maintained in serum-free medium, Neurobasal A supplemented with Glutamax, 2% SM1 Neurocult, and 1% PS. Conditioned medium and tissue lysates were harvested for subsequent biochemical analyses.

### Mouse Studies

#### Animal model

All animal studies were conducted following approval by the Institutional Animal Care and Use Committee and in compliance with NIH guidelines. The CRND8-Tg mice overexpressing amyloid precursor protein (*APP*) begin to develop Thioflavin-S positive Aβ amyloid plaques by 3 months of age [52]. CRND8-Tg mice were bred within the facility, kept in groups of three to five per cage, and provided unrestricted access to food and water under a 12-hour light/dark cycle. Both male and female mice were used. Group sizes were n = 7 per group.

#### Neonatal rAAV injection and brain tissue processing

rAAVs were delivered intracerebroventricularly to neonatal mice on postnatal day 0 (P0) as previously described [53]. Two microliters of rAAV2/8 encoding Sdc4 were injected bilaterally into the cerebral ventricles of CRND8-Tg mice. CRND8-Tg mice receiving PBS injections served as CRND8-Tg controls. Non-transgenic (non-Tg) mice were injected with either rAAV2/8-Sdc4 or PBS. All mice were euthanized 4 months after injection. One hemibrain was fixed overnight in 4% paraformaldehyde at 4 °C, followed by processing and paraffin embedding for immunohistochemical analysis. The contralateral hemibrain was snap-frozen in isopentane on dry ice and stored at −80 °C until it was thawed and homogenized for ELISA quantification of Aβ species and for Western blot analysis.

The hemisphere was cryo-pulverized in liquid nitrogen. Sequential extraction was performed to obtain different solubility fractions previously described [54, 55]. Briefly, homogenates were first extracted in RIPA buffer (50 mM Tris-HCl, 150 mM NaCl, 1% Triton X-100, 0.5% deoxycholate, and 0.1% SDS) with protease inhibitors followed by ultracentrifugation at 100,000 xg for 1 hour at 4 °C to obtain the RIPA-soluble fraction. The resulting pellets were subsequently re-extracted in 2% SDS with protease inhibitors and centrifuged under the same conditions to obtain the SDS-soluble fraction. The remaining insoluble material was further extracted in 70% formic acid (FA) by sonication and centrifugation to generate the FA-soluble fraction. Prior to analysis, FA extracts were neutralized by dilution in Tris-based neutralization buffer.

#### Histological and immunohistochemical analysis of mouse brain sections

Mouse brain immunohistochemistry was performed using paraffin-embedded sections (5 μm). Thioflavin-S (Thio-S) staining was conducted following previously reported protocols [56]. Paraffin sections were subjected to immunohistochemical staining using a biotinylated pan-Aβ antibody (mAb5, 1:500) or an anti-FLAG antibody. Signal detection was carried out with the Vectastain Elite ABC Kit and visualized using the 3,3’-Diaminobenzidine Substrate Kit, followed by hematoxylin counterstaining. Digitized slides were analyzed using the ImageScope software. In brief, at least three sections per sample, at least 30 μm apart, were imaged and plaque burden was quantified.

### Statistical analysis

Statistical analyses were performed as indicated in the corresponding figure legends. For comparisons between two groups, either a paired or an unpaired t-test was used. Multiple t-tests with two-stage step-up correction (Benjamini, Krieger, and Yekutieli) were applied when multiple comparisons were performed between two groups. For comparisons involving more than two groups, statistical significance was assessed using one-way or two-way analysis of variance (ANOVA), followed by Tukey’s post hoc test. A *p* value < 0.05 was considered statistically significant. All analyses were performed using GraphPad Prism version 11.0.0, which was also used for graph generation and data visualization. Data are presented as mean ± standard error of the mean (SEM).

### Use of AI tools

During manuscript preparation, AI-assisted tools were used for language editing and preparation of the graphical abstract. The authors critically reviewed and take full responsibility for the final content.

## Results

### Shared upregulation of syndecan-4 in CRND8-Tg mice and Alzheimer’s disease patients and its association with Aβ

Proteomic studies of human and amyloid-depositing mouse model brains identify multiple HSPGs that are significantly and reproducibly altered in AD and Aβ amyloid mouse models (Fig. 1A) [6, 13, 26, 41]. Though AGRIN has been the most intensively studied HSPG in the context of its role in AD, it is not the most altered in these proteomic studies. SPOCK1-3 and SDC4 are consistently observed to be the most upregulated HSPGs in both AD and Aβ amyloid mouse models, including across other brain regions and in other APP transgenic mice [10, 26, 57, 58]. In 18-month-old CRND8-Tg mice brains, multiple chondroitin sulfate proteoglycans (CSPGs) are significantly decreased, whereas in AD, the changes in CSPGs are more variable. Numerous proteins involved in HS/CS metabolism are also altered in CRND8-Tg mice, fewer of these are detected in humans and typically show smaller changes. Many matrisome proteins, which are among the most changed proteins in human and mouse proteomes, including APOE, Aβ, MDK, PTN, and VTN, are known to bind HSPGs [6]. As we have previously shown, SDC4 to be present in plaques, dystrophic processes around plaques, and to colocalize with a subset of neurofibrillary tangles [58] (Fig. 1B), we focused on SDC4, as it has links to both amyloid and tau pathology, as well as APOE, for which it can serve as a coreceptor.

**Figure 1.**
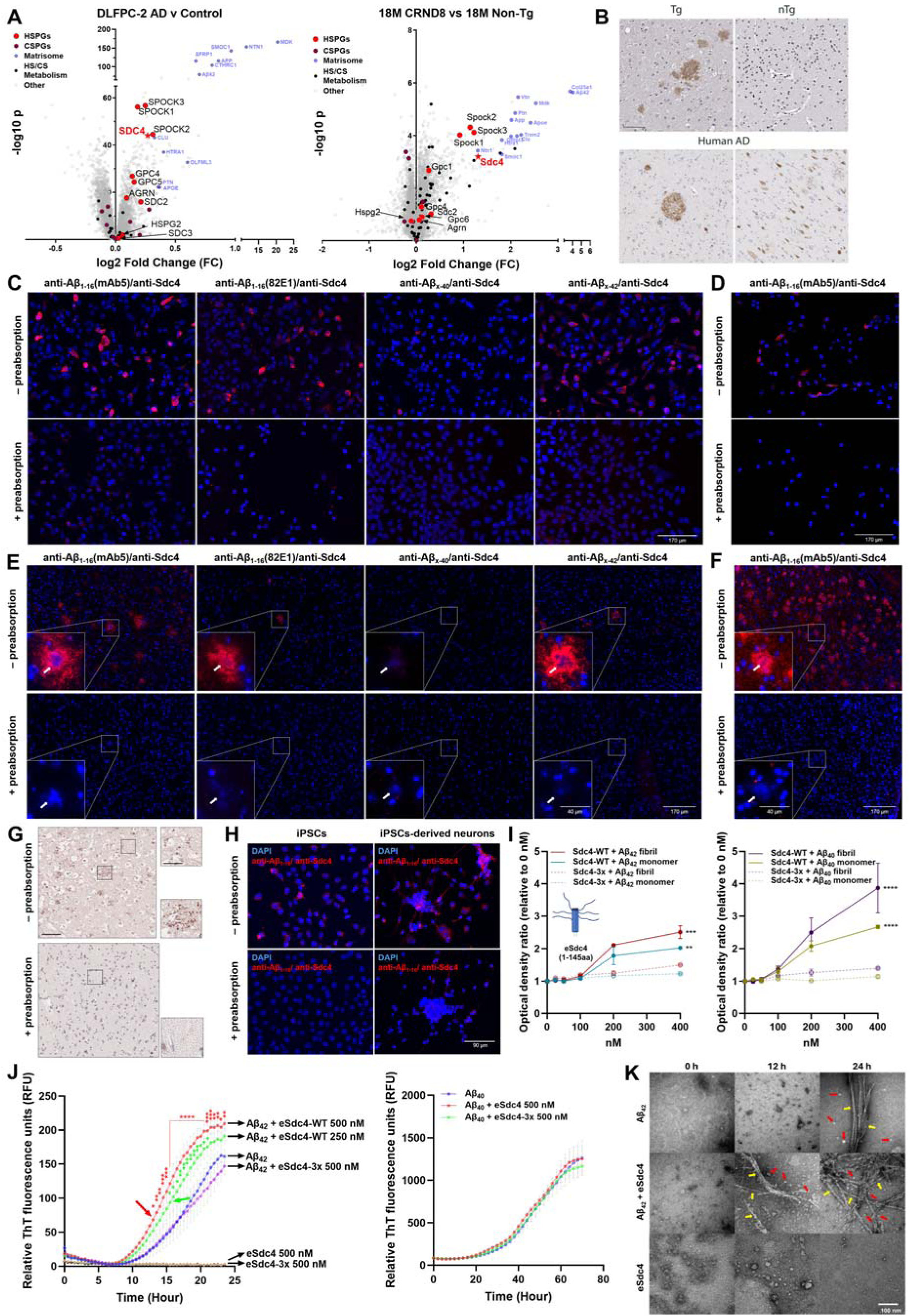
Proteomic convergence-based identification of Sdc4 and its association with Aβ spatial proximity and aggregation. **(A)** Volcano plot of shared differentially expressed proteins (DEPs) between Alzheimer’s disease (AD) patients and healthy controls (Left), and between 18-month-old CRND8-Tg and non-Tg mice (Right). Each point represents a protein, and colors indicate protein categories: red, HSPGs; Tyrian purple, CSPGs; semi-transparent blue, matrisome; black, HS/CS metabolism; white, others. **(B)** Sdc4 immunostaining in AD mouse models and human AD brain. Upper panels: Sdc4 immunoreactivity in brain sections from CRND8-Tg and nTg control mice. Lower panels: Sdc4 immunostaining in human AD brain, with prominent signal in amyloid plaques (left) and neurofibrillary tangles (right). The image with the plaque is from a 79-year-old male (high Alzheimer’s disease neuropathological changes); the section with the tangles is from a 79-year-old female with low Alzheimer’s disease neuropathological changes. Scale bar = 50 μm. **(C)** Proximity ligation assay (PLA) was performed in H4 cells co-expressing amyloid precursor protein containing the Swedish mutation (APP_swe_; K595N/M596L) and Sdc4. Red fluorescence indicates PLA signals, reflecting spatial proximity (<40 nm) between Sdc4 and Aβ. Sdc4 antibody was combined with antibodies recognizing Aβ_1–x_(mAb5), Aβ_1–x_ (82E1), Aβ_x–40_ (13.1.1), or Aβ_x–42_ (2.1.3). The upper panels show PLA signals detected without epitope absorption. The lower panels show corresponding conditions following preabsorption with peptides to competitively mask antibody-recognized Aβ epitopes prior to incubation. Peptides corresponded to the epitopes recognized by each antibody: Aβ_1–16_ peptide for mAb5 and 82E1, Aβ_1–40_ peptide for 13.1.1, and Aβ_1–42_ peptide for 2.1.3. Scale bar = 170 μm. **(D)** PLA analysis in H4 cells expressing BRI-Aβ_42_ and Sdc4. PLA signals (red) were detected using mAb5 and Sdc4 antibodies. Absorption control is shown in the lower panel. Scale bar = 170 μm. **(E)** Brain sections from 12-month-old CRND8-Tg mice were analyzed by PLA using antibody pairs described in (C). Specificity of PLA signals was validated by pre-incubation of antibodies with corresponding peptides. Scale bar = 170 μm; inset = 40 μm. **(F)** PLA in brain sections from 12-month-old BRI-Aβ_42_ mice. Representative images of Sdc4-Aβ proximity signals detected with mAb5 and Sdc4 antibodies. Scale bar = 40 μm. **(G)** Sdc4 and Aβ proximity was detected by brightfield PLA in human AD brain sections. Absorption control is shown. Hematoxylin was used as a nuclear counterstain. Scale bar = 150 μm; inset = 50 μm. **(H)** PLA analysis in hiPSCs and hiPSC-derived neurons. PLA signals (red) were detected using mAb5 and Sdc4 antibodies. The lower panel shows the absorption control following pre-incubation with Aβ_1–16_ peptide. Scale bar = 90 μm. **(I)** Binding curves of Sdc4 measured with increasing concentrations (0–400 nM) using plates coated with Aβ_42_ (left) and Aβ_40_ (right) in monomeric or fibrillar forms. Signal intensities were normalized to the 0 nM condition. n = 2 replicates. **(J)** Thioflavin T fluorescence assay demonstrating Aβ_42_(left) and Aβ_40_ (right) aggregation kinetics in the presence of wild-type eSdc4 (eSdc4-WT) and a heparan sulfate-deficient eSdc4 mutant harboring S44A, S62A, and S64A substitutions (eSdc4-3x) at the indicated concentration. Values represent the average of three time points for Aβ_42_ or six time points for Aβ_40_ over time. n = 6 replicates for Aβ_42_; n = 3 replicates for Aβ_40_. **(K)** Transmission electron microscopy images of Aβ_42_ in the absence or presence of eSdc4-WT collected from an independent aggregation reaction. Red and Yellow arrows indicate Aβ_42_ oligomers and fibrils, respectively. Scale bar, 100 nm. *Nuclei were counterstained with DAPI (blue). *Data are presented as mean ± SEM. *Statistical significance was determined by two-way ANOVA followed by Tukey’s multiple comparisons test. *p < 0.05, **p < 0.01, ***p < 0.001, ****p < 0.0001.

Given the enrichment of SDC4 around amyloid plaques, the proximity of Sdc4 to Aβ was examined using Proximity ligation assay (PLA). To validate the PLA, we first tested the antibody pairs in transiently transfected H4 cells. H4 cells co-expressing APP_swe_ and Sdc4 confirmed that the PLA signals were only produced when APP and Sdc4 were co-transfected (Fig. 1C). Preabsorption with a peptide (Aβ_1-16_) targeting the mAb5 epitope almost entirely reduced the PLA signals, confirming the specificity of the Sdc4-Aβ PLA. (Additional controls for the PLA are shown in Supplementary Fig. 1). End-specific antibodies, including 82E1 (Aβ_1-x_) and 2.1.3 (Aβ_x-42_), also produced strong PLA signals that were reduced by the corresponding peptides, with 13.1.1 (Aβ_x-40_) generating a weaker but detectable signal. Additionally, co-expression of a fusion protein, BRI-Aβ_42_, based on the ITM2B protein (also known as BRI2), that we have previously utilized to generate Aβ_42_ independently from APP [59], showed that co-expression of BRI-Aβ_42_ and Sdc4 also revealed strong PLA signals with the mAb5/Sdc4 antibody pair (Fig. 1D).

Having validated the Aβ-Sdc4 PLA, we evaluated brain tissue sections from 12-month-old CRND8-Tg and BRI-Aβ_42_ mice to examine the proximity of Sdc4-Aβ in AD pathophysiology. Distinct PLA puncta were detected both within and near the plaques, with signals largely eliminated by preabsorption (Fig. 1E, F). Anti-Aβ_1-x_ (82E1)/Sdc4 and anti-Aβ_x-42_ (2.1.3)/Sdc4 pairs confirmed Sdc4 proximity within amyloid plaques, while the anti-Aβ_x-40_ (13.1.1)/Sdc4 pair again yielded only faint signals. Plaque-associated PLA signals from the mAb5/Sdc4 antibody pair were detected in the brain of an AD patient and were blocked by pre-incubation of mAb5 with Aβ_1-16_ peptide (Fig. 1G). Unexpectedly, a strong PLA signal for Aβ-SDC4 was observed in hiPSCs and hiPSC-derived neurons (Fig. 1H). The signal, which was largely localized to the cell and cell membrane, was completely abolished by preabsorption. These data demonstrate that Aβ and Sdc4 are closely associated in human brain cells in a non-pathogenic, physiologic setting.

### The heparan sulfated Sdc4 ectodomain associates with Aβ and impacts Aβ aggregation

To directly investigate interactions between Sdc4-Aβ binding, we purified the wild-type syndecan-4 ectodomain (eSdc4-WT) and a heparan sulfate-deficient mutant (eSdc4-3x) carrying Ser44Ala, Ser62Ala, and Ser64Ala substitutions (Supplementary Fig. 2A). An ELISA-based binding assay using plates coated with monomeric or fibrillar Aβ_40_ or Aβ_42_ showed that eSdc4 bound both Aβ species in a concentration-dependent manner across 0-400 nM (Fig. 1I). eSdc4-WT exhibited significantly higher binding signals than eSdc4-3x, with a tendency toward higher signals on Aβ_40_ compared to Aβ_42_.

The effect of eSdc4 on Aβ_42_ and Aβ_40_ aggregation was then assessed by ThT fluorescence kinetics. eSdc4-WT accelerated Aβ_42_ aggregation as evidenced by a shortened lag phase, while eSdc4-3x produced no measurable effect (Fig. 1J and Supplementary Fig. 2B). In contrast, neither construct altered Aβ_40_ aggregation kinetics. Transmission electron microscopy revealed that fibrils were detectable at 12 hours only in the presence of eSdc4-WT, whereas Aβ_42_ alone contained no fibrils (Fig. 1K). By 24 hours, fibrils were observed under both conditions. Fibrils formed in the presence of eSdc4-WT were thicker and exhibited a dense, uneven appearance. In addition, numerous small, rounded oligomers were observed in the eSdc4-WT condition compared to Aβ_42_ alone.

### Syndecan-4 reduces Aβ levels by inhibiting APP processing

To explore the broader hypothesis that proteins within the amyloid responsome (i.e., proteins that accumulate in response to amyloid) might not just be markers of disease but play a disease-modifying role, we evaluated whether Sdc4 impacted APP processing and Aβ production. Transient co-expression of APP_swe_ with Sdc4 consistently and dramatically reduced secreted Aβ levels relative to GFP-expressing controls in both HEK293T and CHO cells (Fig. 2A, B). Western blot analysis of these cells revealed pronounced decreases in soluble APPα (sAPPα), C83, and C99 in Sdc4 co-expressing cells. In these studies, soluble APPβ (sAPPβ) was not detectable (Fig. 2C, D). Dose-response studies showed that increasing Sdc4 expression led to progressively lower levels of Aβ_1-x_ (Fig. 2E). We next evaluated the impacts of Sdc4 on APP expression in both pooled stable and clonal stable lines. Both the APP_695_WT + Sdc4 pooled and APP_695_WT + Sdc4 single clonal cells showed significantly reduced secreted Aβ levels compared with 2B7 cells, which express only APP_695_WT (Fig. 2F). Sdc4 also markedly decreased sAPPα and sAPPβ levels, while C99 was not detected (Fig. 2G, H). C83 appeared to be decreased in the cells expressing Sdc4, but these changes did not reach statistical significance. MALDI-TOF mass spectrometry (MS) of Aβ immunoprecipitated (IP) from conditioned medium further confirmed Sdc4-induced reductions in secreted Aβ levels (Fig. 2I, J). These IP-MS studies also did not confirm the larger reduction in Aβ_x-42_ observed by ELISA, indicating that all species of Aβ were decreased equally.

**Figure 2.**
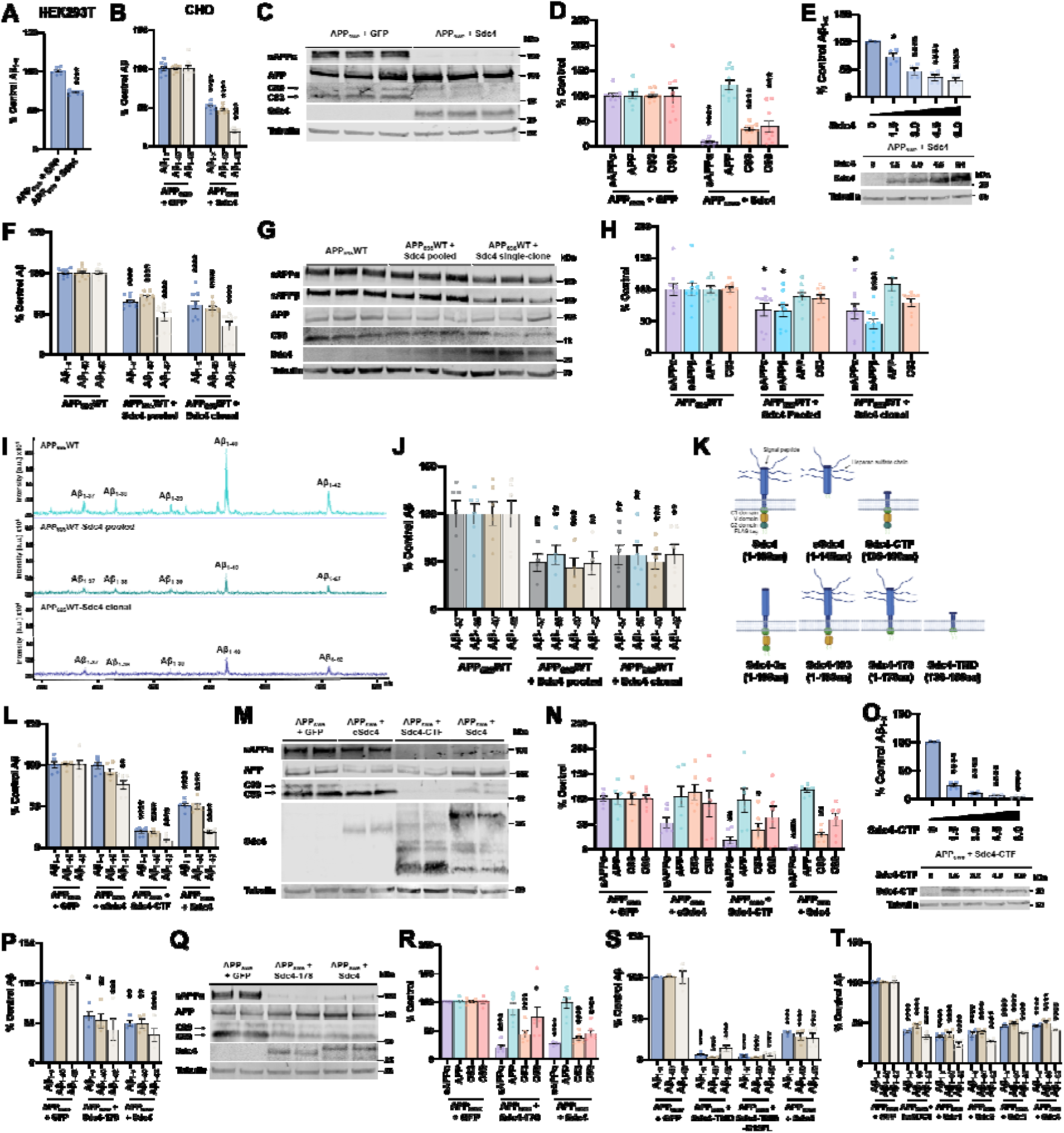
Sdc4 alters APP processing and. **A**β **levels in cells. (A)** ELISA quantification of secreted Aβ species from HEK293T cells transiently co-transfected with amyloid precursor protein with K595N/M596L substitutions (APP_swe_) and syndecan-4 (Sdc4). n = 9 replicates. **(B)** ELISA quantification of secreted Aβ species from CHO cells transiently co-transfected with APP_swe_ and Sdc4. GFP was used as a control. n = 8 replicates. **(C)** Representative Western blot of APP, APP C-terminal fragments (APP-CTFs), and soluble APPα (sAPPα). **(D)** Quantification of the immunoblots shown in (C). n = 9 replicates. **(E)** Dose-dependent effect of Sdc4 on secreted Aβ species. Cells were transfected with increasing amounts of Sdc4 plasmids (0 to 6.0 units). A uniform total amount of transfected DNA was maintaine across all conditions by adding an empty vector to compensate for differences in plasmid amounts. Results are normalized to the control (0) and expressed as % Control. Statistical significance was determined by comparing it against the control (0). n = 4 replicates. **(F)** Levels of Aβ species secreted in stable cells expressing APP_695_WT alone or co-expressing syndecan-4 (Sdc4) in pooled populations and an isolated single-cell clone. n = 9 replicates. **(G)** Representative Western blot of APP and its CTFs in stable cell line lysates, and soluble APP fragments (sAPPα and sAPPβ) in the corresponding conditioned medium. **(H)** Quantitative analysis of immunoblots presented in (G). n = 9 replicates. **(I)** Representative MALDI-TOF mass spectra of secreted Aβ peptides from stable CHO cells. Characteristic peaks corresponding to Aβ species are indicated. **(J)** Quantitative analysis of the MALDI-TOF mass spectrometry data presented in (I). n = 6 replicates for APP_695_WT and APP_695_WT + Sdc4 clonal. n = 4 replicates for APP_695_WT + Sdc4 pooled. **(K)** Schematic of Sdc4 domain structure and variants. **(L)** ELISA quantification of secreted Aβ species from APP_swe_ co-expressed with ectodomain Sdc4 (eSdc4) or Sdc4 C-terminal fragment (Sdc4-CTF). n = 6 replicates. **(M)** Corresponding Western blots of APP, sAPPα, APP-CTFs, and Sdc4 variants expression. **(N)** Relative protein levels derived from immunoblots shown in (M). n = 6 replicates. **(O)** Gradient analysis of Sdc4-CTF expression. Cells were transfected with graded amounts of Sdc4-CTF plasmid (0 - 6.0 units). To ensure equal total DNA input across conditions, an empty vector was added as needed to balance plasmid quantities. Data were normalized to the 0 condition and are presented as a percentage of control. Statistical comparisons were performed relative to the 0 condition. n = 4 replicates. **(P)** Aβ species levels by truncated Sdc4 containing amino acids 1 to 178 (Sdc4-178). n = 4 replicates. **(Q)** Representative Western blot figure of APP processing components. **(R)** Signal quantification from immunoblots in (Q). n = 4 replicates. **(S)** Effect of the Sdc4 transmembrane domain (136–156 aa; Sdc4-TMD) on Aβ_1-x_ levels. The Gly157Leu mutant construct (Sdc4-TMD-G157L) was also tested. n = 4 replicates. **(T)** Human SDC4 and other syndecan family members, including murine Sdc1, Sdc2, and Sdc3, were examined for their effects on Aβ levels from cells co-transfected with APP_swe_. n = 4 replicates. *All data are expressed as a percentage of the control group. *Data are presented as mean ± SEM. *Statistical significance was determined by two-way ANOVA followed by Tukey’s multiple comparisons test. *An unpaired Student’s *t*-test was used for (A). *Statistical comparisons were performed using multiple t-tests for (B) and (D). *Data were analyzed by one-way ANOVA followed by Tukey’s multiple comparisons test for (E) and (O). *APP and CTFs quantifications normalized to beta-tubulin. *p < 0.05, **p < 0.01, ***p < 0.001, ****p < 0.0001.

### Syndecan-4 impacts on Aβ levels and APP processing are mediated by its transmembrane domain (TMD)

We conducted structure-function studies to identify the domains of Sdc4 that mediate impact on APP and Aβ. We generated mutants of Sdc4 based on its functional domain organization as illustrated in Fig. 2K. We first tested the impacts of the ectodomain only (eSdc4) that contains the HS chains, and the carboxyl-terminal fragment (CTF) of Sdc4 that is like what would be generated following ectodomain shedding. Expression of Sdc4-CTF with APP_swe_ in CHO cells reduced Aβ levels more effectively than full-length Sdc4, whereas eSdc4 had minimal impact, causing only a slight reduction in Aβ_42_ (Fig. 2L). Western blots showed reductions in sAPPα and C83 in cells expressing Sdc4-CTF (Fig. 2M, N). As with intact Sdc4, Aβ levels decreased progressively with increasing amounts of Sdc4-CTF (Fig. 2O). Full-length Sdc4-3x that lacks HS chains (like the eSdc4-3x mutant) reduced Aβ comparably to Sdc4 (Supplementary Fig. 3B). Analysis of C-terminal truncated Sdc4 variants showed that Sdc4-193 (devoid of the PDZ-containing C2 domain) and Sdc4-178 (lacking both the V and C2 domains) reduced Aβ levels and APP cleavage similarly to full-length Sdc4 (Fig. 2P-R, and Supplementary Fig. 3C). Collectively, these data pointed to the TMD of Sdc4 as a mediator of the effect of Aβ and APP. We tested this assertion directly by expressing the TMD of Sdc4 (Sdc4-TMD) along with a few residues at the cytoplasmic membrane face (136–156 aa). The Sdc4-TMD reduced Aβ_1-X_ levels, as did an additional Sdc4-TMD mutant (Sdc4-TMD-G157L) designed to disrupt glycine zipper-mediated homodimerization (Fig. 2S and Supplementary Fig. 3D).

Sequence alignment revealed that the TMDs of the larger syndecans are highly conserved (Supplementary Fig. 3F). Thus, we assessed whether other syndecan family members also modulate Aβ production. Cells were co-transfected with APP_swe_ and GFP or human syndecan-4 (huSDC4) and mouse Sdc1, Sdc2, Sdc3, and Sdc4. All syndecans significantly decreased Aβ levels similarly to Sdc4 (Fig. 2T and Supplementary Fig. 3E). As the TMD of these proteins is highly conserved, these studies further support the syndecan TMD as a mediator of these effects on APP and Aβ.

### Syndecan-4 co-localizes with APP and Aβ in cells co-expressing both proteins

The impacts of Sdc4 on APP processing and Aβ were not consistent with inhibition of α-β-or γ-secretases but instead suggested a broader alteration in APP trafficking. Thus, we evaluated the subcellular distribution of Sdc4 and APP in H4 cells transiently co-transfected using APP-GFP with Sdc4, eSdc4, or Sdc4-CTF. Immunofluorescence revealed co-localization of Sdc4 with APP-GFP within large vesicular structures (Fig. 3A). Sdc4-CTF also co-localized with APP-GFP in vesicles, whereas eSdc4 showed no co-localization and did not induce the formation of large vesicular structures. These large vesicles were positive for the autophagy markers ATG5 and LC3B, as well as cathepsin D and flotillin in a subset of vesicles (Fig. 3B). They were negative for other markers, such as EEA1, LAMP1, and Rab family proteins (Supplementary Fig. 5B, C), with no detectable co-localization. In CHO stable cell lines expressing APP_695_WT with Sdc4 (single clones), Sdc4 co-localized with both Aβ_1–x_ and the N-terminal APP, although large vesicles were not observed (Supplementary Fig. 5A).

**Figure 3.**
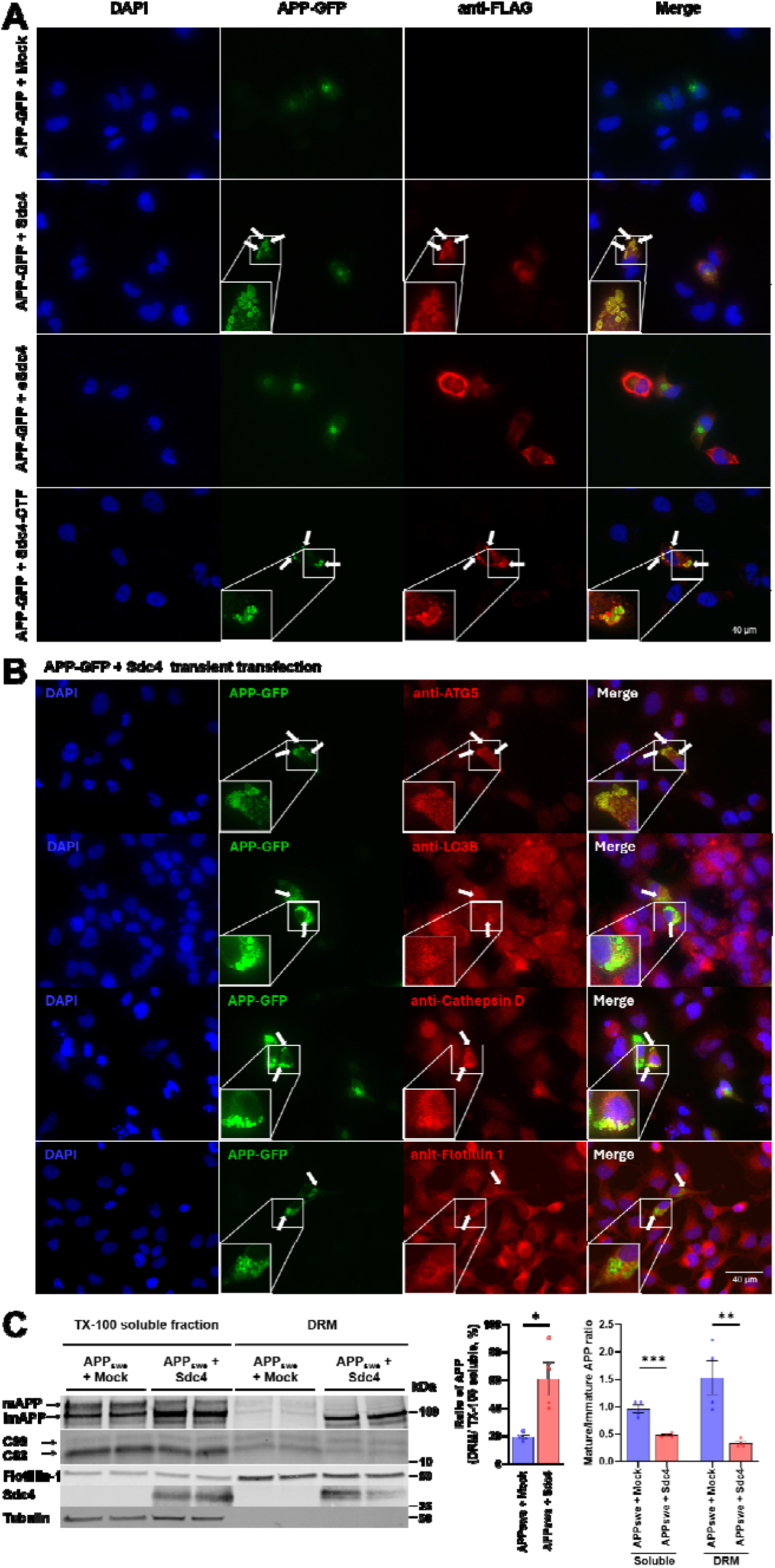
Co-localization of APP and Sdc4 in vesicles positive for autophagy–lysosomal trafficking network proteins. **(A)** Subcellular localization of APP-GFP and Sdc4 variants in H4 cells co-transfected with APP-GFP and full-length syndecan-4 (Sdc4), the ectodomain of syndecan-4 (eSdc4), or the C-terminal fragment of syndecan-4 (Sdc4-CTF). Mock represents an empty vector. Green fluorescence corresponds to APP-GFP, and red fluorescence indicates Sdc4 variants. Sdc4 variants were visualized with anti-FLAG antibody. **(B)** Distribution of APP-GFP and autophagy-lysosomal pathway markers, including autophagy-related 5 (ATG5), LC3B, and cathepsin D, in H4 cells following co-transfection with APP-GFP and Sdc4. Green fluorescence for APP-GFP and red fluorescence for each marker. **(C)** Sdc4-induced re-localization of APP into lipid rafts in HEK293T cells. The ratios of detergent-resistant membrane (DRM) to Triton X-100 (TX-100) soluble APP, as well as the ratio of mature to immature APP, were calculated and plotted. n = 4 replicates. * Nuclei were stained with DAPI (blue). * Co-localization is indicated by white arrows. * Scale bar = 40 μm.

APP distribution between detergent-soluble and detergent-resistant membrane fractions was examined in HEK293T cells co-expressing APP_swe_ and Sdc4 (Fig. 3C). The DRM fraction was enriched in flotillin-1 and devoid of tubulin, confirming its identity as a raft-enriched membrane fraction. Sdc4 overexpression increased the proportion of APP in the DRM fraction relative to the TX-100 soluble fraction. Within each fraction, Sdc4 lowered the mature/immature APP ratio compared with the mock control, in both the soluble and DRM fractions. Notably, whereas the mock control showed a higher mature/immature ratio in the DRM fraction than in the soluble fraction, Sdc4 did not exhibit this increase, owing to a pronounced accumulation of immature APP in the DRM fraction.

### Overexpression of Sdc4 reduces amyloid pathology in neuronal and mouse model systems

ICV delivery of rAAV2/8-CBA-Sdc4 to P0 mice resulted in robust Sdc4 expression in the brains of CRND8-Tg and nTg mice (Fig. 4A), confirming successful viral transduction and protein expression. Overexpression of Sdc4 prevented accumulation of Aβ-positive plaques and resulted in reduced total amyloid burden in 4-month-old Sdc4-expressing CRND8-Tg mice compared with controls (Fig. 4B). Quantification of plaque counts demonstrated a significant 64% decrease in total plaques. ELISA analysis of sequentially extracted brains showed that Sdc4 significantly lowered levels of Aβ_42_ and Aβ_40_ from SDS-(67% and 68% of control) and of Aβ_42_ from FA-extracted fractions (58% of control) in Sdc4-treated mice (Fig. 4C). These data indicate that Sdc4, instead of promoting amyloid deposition *in vivo*, attenuates it, suggesting that the effects on APP processing may override possible impacts of HS-mediated acceleration of amyloid formation. The reductions in Aβ_42_ levels in the RIPA fraction are consistent with this trend but did not reach statistical significance.

**Figure 4.**
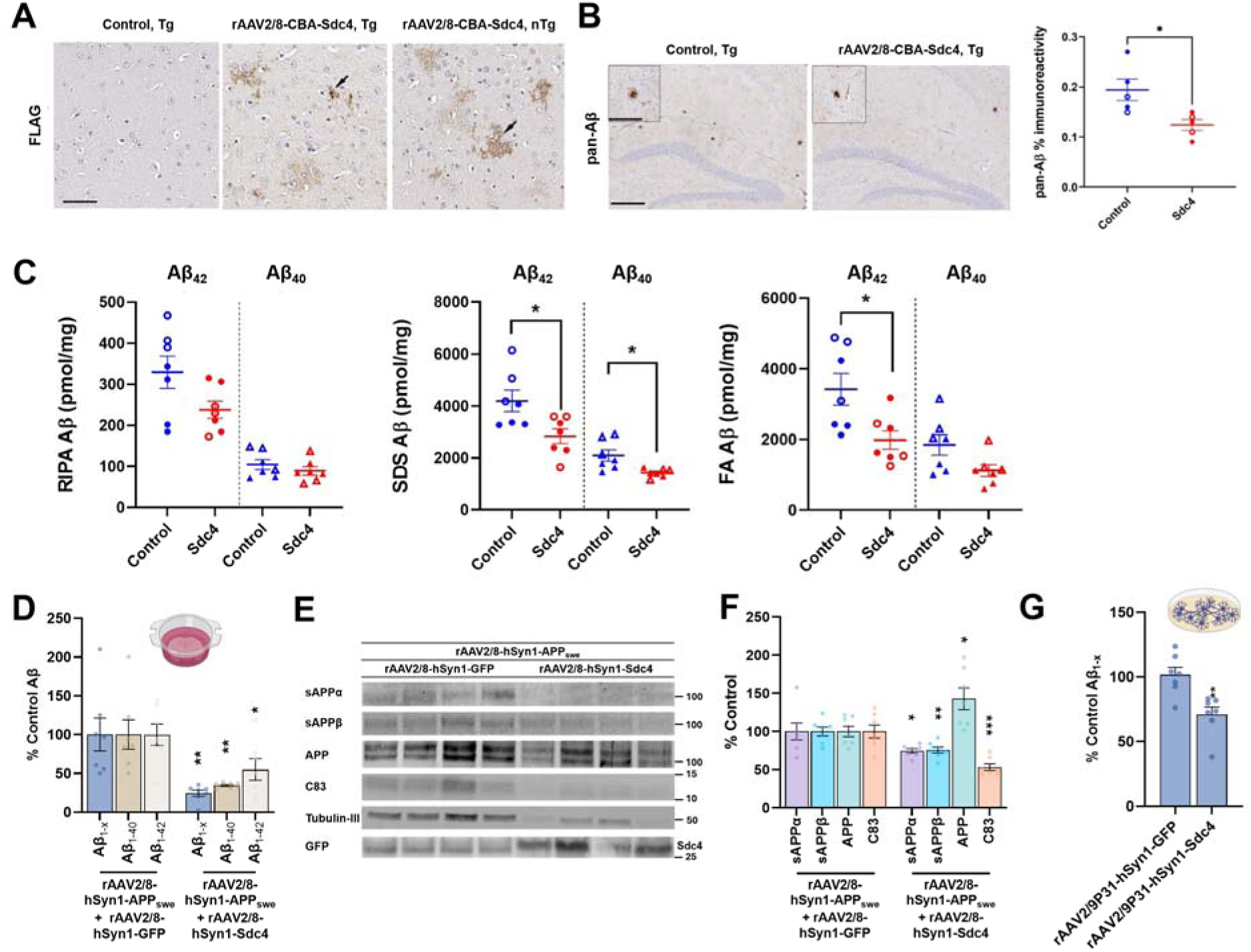
Sdc4 modulation of amyloid deposition in CRND8-Tg mice, including. **A**β **regulation in hiPSC-derived forebrain neurons and mouse brain slice cultures (A)** P0 CRND8-Tg mice were injected ICV with rAAV2/8-CBA-Sdc4 (10^^13^ genomes/mL, 2 μL per ventricle) and brains were harvested at 4 months of age. Overexpression of syndecan-4 (Sdc4) in parenchyma as well as in dystrophic neurons surrounding plaques was visualized by anti-FLAG staining. **(B)** Representative brain sections showing pan-Aβ staining in the cortex and hippocampus of Tg mice. Scale: 250 μm, inset: 50 μm. Amyloid burden in three non-consecutive sections is represented by a scatter dot plot of male (closed circle) and female (open circle). **(C)** RIPA, 2% SDS, and 70% formic acid (FA) extracted Aβ_42_ and Aβ_40_ levels were detected by ELISA and plotted as a scatter dot plot of male (closed circle/triangle) and female (open circle/triangle). **(D)** Levels of secreted Aβ species in the medium of organotypic mouse brain slice cultures transduced by rAAV2/8-hSyn1-APP_swe_ and either rAAV2/8-hSyn1-GFP or rAAV2/8-hSyn1-Sdc4. **(E)** Western blot analysis of APP and its proteolytic fragments (sAPPα, sAPPβ, and C83) in lysates and medium from the brain slice cultures described in (D). **(F)** Quantitative analysis of the immunoblots shown in (E). **(G)** ELISA quantification of secreted total Aβ levels in the conditioned medium from human induced pluripotent stem cells derived neurons (hiPSC-derived neurons) overexpressing GFP or Sdc4. *Data are presented as mean ± SEM. *n = 7 replicates for all experiments. *Statistical comparisons were performed using Paired *t*-test (B), (C), and (G). Multiple t-tests were used for (D) and (F). *p < 0.05, **p < 0.01, ***p < 0.001.

To further evaluate potential impacts of Sdc4 on Aβ production, we examined Sdc4 overexpression in *ex vivo* mouse OBSCs and hiPSC-derived forebrain neurons under conditions of AAV-mediated expression. To confirm AAV functionality in the OBSC system, mouse brain slices were first transduced with rAAV2/8-hSyn1-APP_swe_, resulting in increased Aβ levels and confirming APP overexpression (Supplementary Fig. 7C, D). Subsequently, OBSCs were co-transduced with rAAV2/8-hSyn1-APP_swe_ and rAAV2/8-hSyn1-Sdc4, which led to decreased Aβ levels compared to rAAV2/8-hSyn1-GFP (Fig. 4D). Consistent with cell-based findings, reduced levels of APP proteolytic fragments, including sAPPα, sAPPβ, and C83, were also observed in slice lysates, whereas full-length APP levels were increased (Fig. 4E, F). These data confirmed that, in brain cells where Aβ is not depositing, Sdc4 inhibits Aβ production through impacts on APP processing. Finally, in hiPSC-derived neurons, AAV-mediated expression of Sdc4 decreased Aβ levels in conditioned medium relative to control (Fig. 4G, Supplementary Fig. 7E). For hiPSC-derived neurons expressing endogenous APP, Aβ_40_ and Aβ_42_ levels were below the limit of quantification. These data demonstrate that increased levels of Sdc4 can alter endogenous human neuronal APP processing and decrease Aβ production.

## Discussion

Sdc4 is the most significantly increased transmembrane HSPG in the AD brain. Our findings reveal that different domains of Sdc4 may exert opposing effects on amyloid pathology. The extracellular domain of Sdc4 promotes Aβ_42_ aggregation through its HS chains, whereas full-length Sdc4 suppresses Aβ production by inhibiting APP processing. This functional dichotomy suggests that the overall impact of Sdc4 on amyloid pathology is determined by the balance between its extracellular, HS-dependent, and intracellular, HS-independent, activities. While the aggregation-promoting effects of HS are well established, the ability of Sdc4 to suppress APP processing was unexpected. Importantly, our *in vivo* data indicate that, at least during early stages of amyloid deposition, the inhibitory effects on Aβ production predominate over its aggregation-promoting properties.

Our PLA studies extend previous observations of Sdc4-Aβ co-localization by demonstrating close spatial proximity between the two proteins in both human AD and amyloid-depositing mouse brain tissue. PLA signals were concentrated within and surrounding amyloid plaques, supporting a close physical association between Sdc4 and Aβ *in vivo*. Similar signals were also detected in hiPSCs and hiPSC-derived neurons without overexpression, suggesting that Sdc4-Aβ interactions occur under physiological conditions and may be amplified during disease. Whether this relationship reflects direct binding or co-localization remains to be determined.

Consistent with the established role of HS in amyloidoses [44, 60], the soluble ectodomain of Sdc4 accelerated Aβ_42_ aggregation, whereas an HS-deficient mutant had no effect, identifying HS chains as the critical mediator of this activity. Interestingly, Sdc4 selectively enhanced Aβ_42_ but not Aβ_40_ aggregation, suggesting that specific HS structural features, such as sulfation patterns, may influence this interaction. The altered morphology of Aβ_42_ fibrils formed in the presence of Sdc4 further supports an association between Sdc4 and Aβ aggregates.

In contrast, Sdc4 consistently suppressed APP processing and Aβ secretion across transient and stable expression systems, including OBSCs and hiPSC-derived neurons. Reductions in sAPPα, sAPPβ, C83, and C99, together with altered APP localization, indicate that Sdc4 broadly regulates APP trafficking rather than individual secretase activities. Mechanistically, Sdc4 promoted APP accumulation within flotillin-positive, LC3-and ATG5-associated vesicular structures that also contained active cathepsin D. The absence of markers for early endosomes and mature lysosomes suggests that these compartments represent an intermediate autophagy-related trafficking pathway. These observations support a model in which Sdc4 redirects APP away from amyloidogenic processing and toward autophagy-associated compartments. The association of Sdc4-APP vesicles with flotillin-1 further implicates lipid raft-dependent trafficking in this process. Although amyloidogenic processing has traditionally been linked to cholesterol-rich lipid raft domains [61], our findings suggest that Sdc4 may segregate APP into specialized raft-associated compartments that limit APP-secretase interactions or redirect APP toward degradative pathways. Increased localization of APP within detergent-resistant membrane fractions and flotillin-positive vesicles, together with reduced APP processing, supports this interpretation.

Domain-mapping studies identified the TMD of Sdc4 as necessary for the suppression of APP processing. Given the high evolutionary conservation of the Sdc4 TMD, this finding suggests a previously unrecognized role for the TMD in regulating membrane trafficking. Disruption of the proposed glycine zipper motif and TMD dimerization did not alter Sdc4 activity, indicating that the mechanism is unlikely to depend on these structural features. Similarly, despite extensive ectodomain shedding, we found little evidence that the Sdc4 C-terminal fragment undergoes γ-secretase processing (Supplementary Fig. 4). Instead, the higher apparent molecular weight of TMD-containing fragments suggests the possibility of stable multimeric complexes. These findings parallel reports of non-canonical signaling functions mediated by the Notch1 transmembrane domain [62] and raise the possibility that the Sdc4 TMD serves as an active determinant of membrane organization and cargo trafficking rather than simply a membrane anchor.

Sdc4 exerts a robust inhibitory effect on Aβ secretion and accumulation. In CRND8-Tg mice, Sdc4 overexpression reduced amyloid plaque burden and decreased both Aβ_40_ and Aβ_42_ levels, indicating that despite its dual functions, the net effect of Sdc4 is attenuation of amyloid pathology. Consistent with this finding, PLA revealed extensive Sdc4-Aβ associations *in vivo*, with signals enriched within amyloid plaques and perinuclear compartments (Supplementary Fig. 6), suggesting both extracellular interactions with deposited Aβ and intracellular interactions that may facilitate Aβ sequestration and trafficking. Reduction in Aβ secretion was reproduced in organotypic brain slice cultures and hiPSC-derived neurons, supporting the robustness and translational relevance of this effect. The consistent reduction of Aβ across cellular and tissue models further supports a cell-autonomous role for Sdc4 in regulating APP processing. A useful parallel is SORLA, an endocytic sorting receptor that limits amyloidogenic APP processing by regulating intracellular trafficking [63]. Although the specific compartments involved may differ, our findings suggest that Sdc4 similarly influences APP trafficking, thereby restricting APP access to secretase-enriched compartments and reducing Aβ generation.

These and our previous studies raise numerous questions regarding the role of Sdc4 and other Sdcs in AD. Why is Sdc4 increased in AD relative to Sdc1-3? Sdc4 is thought to be selectively expressed in astrocytes, then why does immunostaining clearly show its presence in neuronal cells in the AD brain, both with and without tangles? Will longer-term *in vivo* studies in amyloid depositing mice show different impacts? Can *in vivo* studies using domain-specific constructs (e.g., overexpression of eSdc4, a HS-deficient, mutant or the CTF) provide further insight into the opposing effects on amyloid formation? Disruption of endosomal and autophagic pathways is widely reported in the AD brain [64–67]. Does increased Sdc4 contribute to those disruptions? Do additional functions of Sdc4 including possible impacts on Aβ and tau clearance or APOE internalization, play a pathophysiologic role in AD? Do other understudied HSPGs such as the SPOCKs contribute?

Several limitations should also be considered. Our PLA studies provide evidence for close association of Sdc4 and Aβ, but it is unclear whether this is simple proximity or actual binding. The data in hiPSCs and hiPSC-derived neurons are especially intriguing to consider in this context as it suggests Sdc4 and Aβ either in proximity (e.g., a similar membrane subdomain) or binding each other in a physiological setting. Our studies have largely relied on overexpression as Sdc4 is increased in the AD brain and mouse models of amyloid deposition. It is unclear what a knockout study might show, especially given the functional overlaps of the syndecan family.

Collectively, these findings identify Sdc4 as a multifunctional regulator of amyloid pathology that acts through distinct and dissectible mechanisms. While the HS chains of Sdc4 promote aggregation, its transmembrane and intracellular domains suppress APP processing, apparently by directing APP to flotillin-positive, autophagy-associated compartments. Across all experimental systems examined, the dominant outcome was reduced Aβ accumulation and amyloid burden, possibly positioning Sdc4 as a net-protective modulator of AD pathogenesis at least during early stages of disease. Importantly, these data provide another example of how proteins in the amyloid responsome appear to not just to be markers of disease but act as potential modifiers of pathological processes, thus providing additional support for the amyloid scaffold hypothesis [6].

## Abbreviations

Aβ: Amyloid-beta
AD: Alzheimer’s disease
APP: Amyloid precursor protein
APP_swe_: Amyloid precursor protein containing the Swedish mutation (K595N/M596L)
ATG5: Autophagy-related protein 5
CSPG: Chondroitin sulfate proteoglycan
CTF: C-terminal fragment
DIV: Days in vitro
DRM: Detergent-resistant membrane
ECM: Extracellular matrix
ELISA: Enzyme-linked immunosorbent assay
eSdc4: Syndecan-4 ectodomain
FA: Formic acid
GAG: Glycosaminoglycan
HSPG: Heparan sulfate proteoglycan
HS: Heparan sulfate
hiPSC: Human induced pluripotent stem cell
ICV: Intracerebroventricular
IP: Immunoprecipitation
MS: Mass spectrometry
NFT: Neurofibrillary tangle
NPC: Neural progenitor cell
OBSC: Organotypic brain slice culture
PLA: Proximity ligation assay
RIPA: Radioimmunoprecipitation assay
Sdc4: mouse syndecan-4
SDC4: Human syndecan-4
SEM: Standard error of the mean
TEM: Transmission electron microscopy
ThT: Thioflavin T
Thio-S: Thioflavin S
TMD: Transmembrane domain
TX-100: Triton X-100
WT: Wild type

## Acknowledgements

We are grateful to the patients and their families for their invaluable contributions to Alzheimer’s disease research.

The UF Neuromedicine Human Brain and Tissue Bank (HBTB) is supported by the Evelyn F. and William L. McKnight Brain Institute, the Center for Translational Research in Neurodegenerative Diseases and the 1Florida ADRC (P30 AG047266). S.P. is supported by the Charlotte and Howard Zimmerman rising star professorship at the Norman Fixel Institute for Neurological diseases.

## Author Contributions

K.H.S. designed and performed experiments, analyzed data, and wrote the manuscript. Y.R. and C.M. assisted and conducted cell-based experiments and protein purification. P.A. and F.A. cultured hiPSCs and hiPSC-derived neurons and conducted PLA experiments. D.R. produced rAAVs. B.D.M., Y.Y., X.L., and M.B. performed OBSC and *in vivo*-related experiments. N.T.S. conducted proteomics and related analyses. W.T. participated in scientific discussions and provided intellectual input. L.L. conducted inter-laboratory ELISA validation. M.E.P. and B.R.R. performed MALDI-TOF analyses. S.P. conducted experiments using human samples. Y.L. and T.E.G. contributed to study design, data interpretation, and manuscript revision, supervised the study, and secured funding. All authors contributed to manuscript editing and approved the final manuscript.

## Data Availability

The datasets generated and/or analyzed during the current study are available from the corresponding author upon reasonable request. All data supporting the findings of this study are included in the article and its supplementary information files.

## Declarations

### Ethics Approval and Consent to Participate

All animal procedures were approved by the Institutional Animal Care and Use Committee (IACUC) of Emory University and were conducted in accordance with the National Institutes of Health Guide for the Care and Use of Laboratory Animals. Human postmortem brain tissues were obtained from established brain repositories under protocols approved by the respective institutional review boards. The use of de-identified human specimens was conducted in accordance with applicable ethical guidelines and regulations.

### Consent for Publication

Not applicable.

