## supplemental files for "Syndecan-4 exerts canonical heparan sulfate-dependent and noncanonical heparan sulfate-independent functions that regulate Aβ amyloid homeostasis"

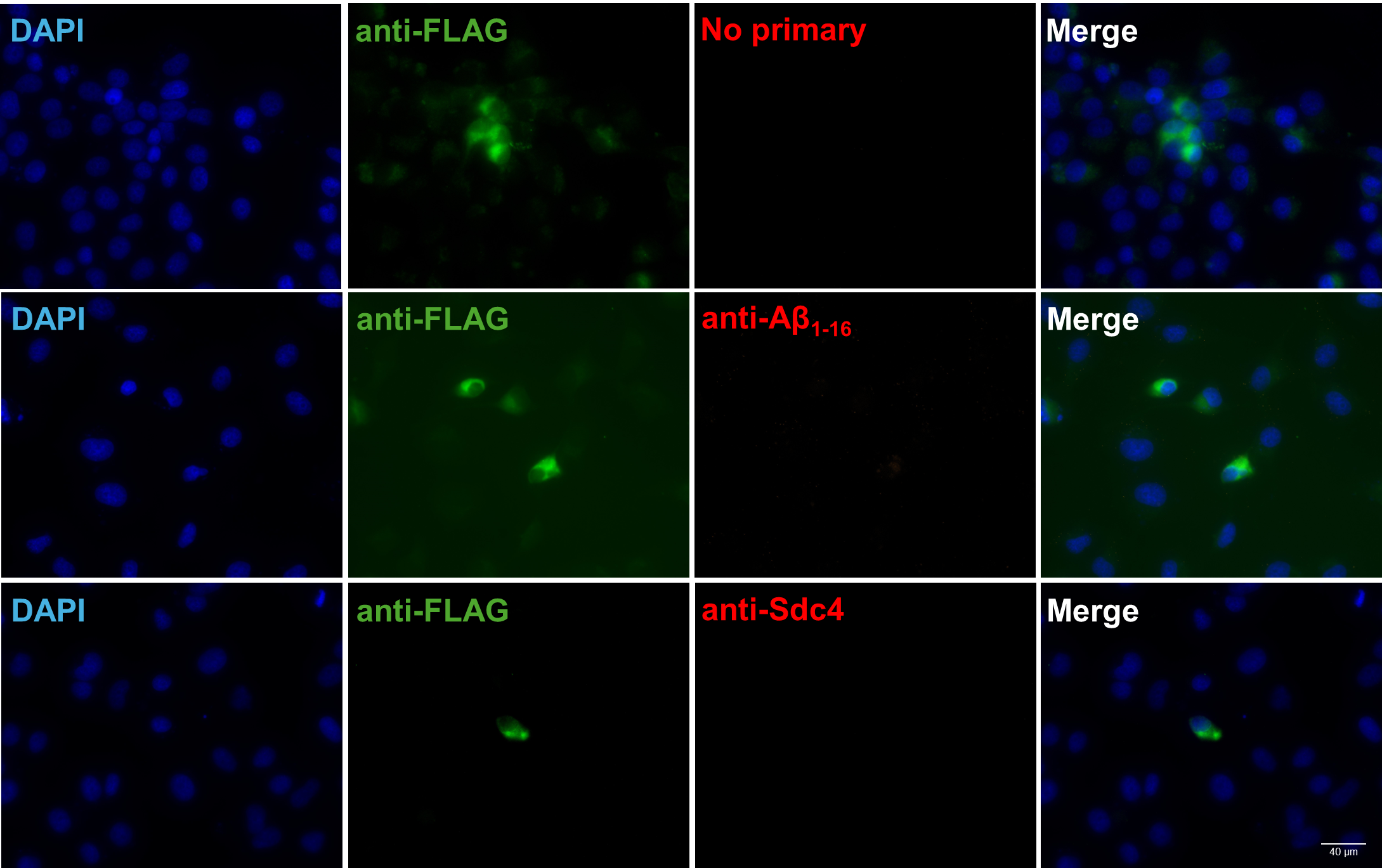

**Supplementary figure 1. Proximity ligation assay (PLA) control conditions.** PLA control conditions included no primary antibody or individual primary antibodies alone targeting Aβ_1–16_ (mAb5) and the ectodomain of syndecan-4 (eSdc4) to evaluate assay specificity. Green displays overexpression of Sdc4 detected by the FLAG antibody.
*Nuclei were counterstained with DAPI (blue).
*Scale bar = 90 μm.

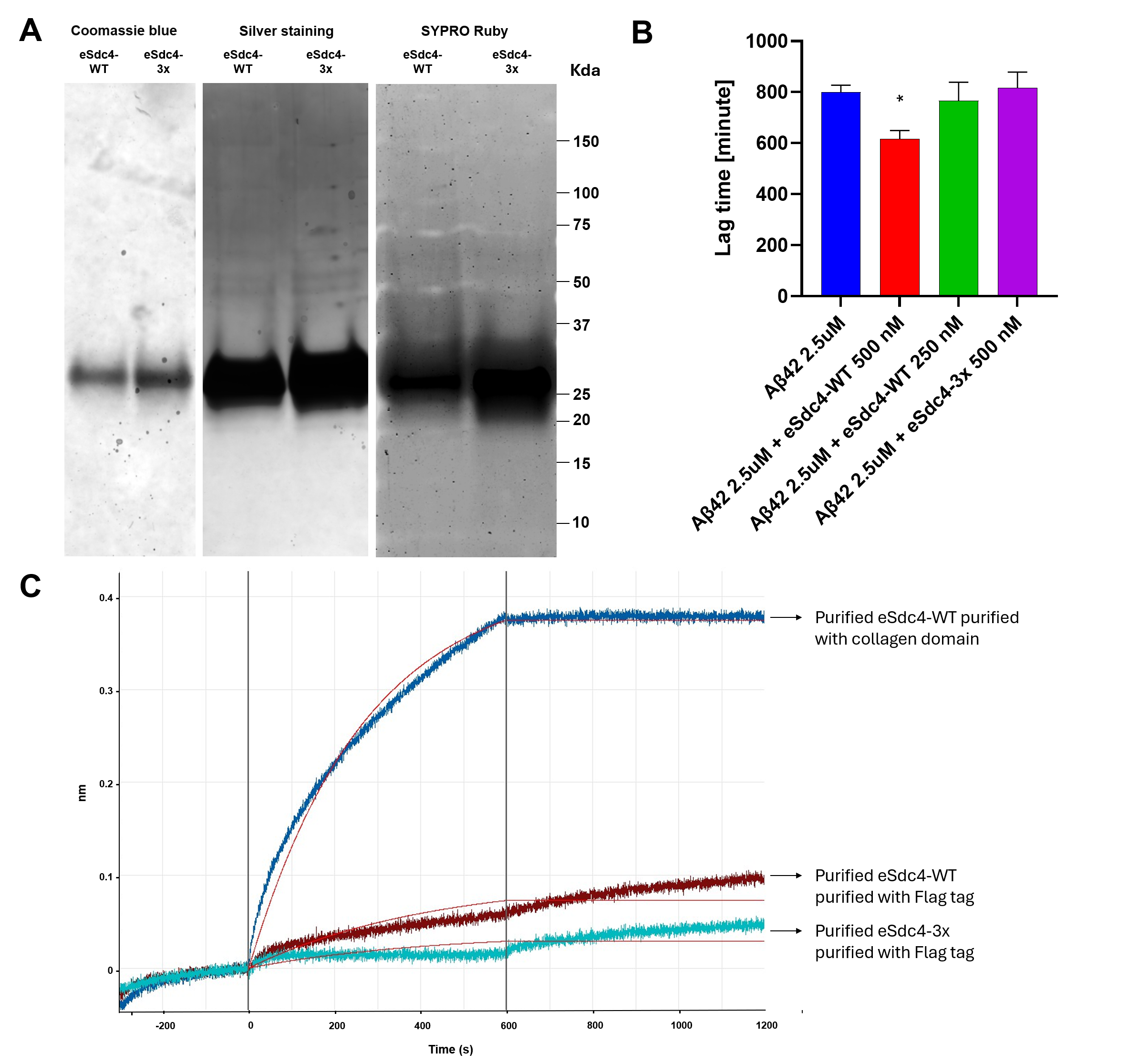

**Supplementary figure 2. Coomassie blue staining of purified syndecan-4 ectodomain proteins and quantification of Aβ_42_ aggregation lag time. (A)** Coomassie blue staining of purified wild-type (WT) syndecan-4 (Sdc4) and a heparan sulfate–deficient Sdc4 ectodomain mutant harboring S44A, S62A, and S64A substitutions (eSdc4-3x), expressed in HEK293T cells. **(B)** Lag times derived from Figure 3A were quantified and presented as bar graphs. Lag time was calculated as the interval between reaction start and the point at which the fluorescence signal first exceeded the threshold. The threshold was defined as the mean of the lowest 10 fluorescence values during the baseline period plus ten times the standard deviation of those baseline values. **(C)** Biolayer interferometry (BLI) binding kinetics were measured using purified eSdc4-WT and eSdc4-3x purified with Flag antibody. Additionally, the eSdc4-WT construct containing the collagen domain was expressed and purified by nickel affinity chromatography, processed to remove the collagen domain, and subjected to binding kinetic analysis. Sensorgrams display association and dissociation phases following exposure to Aβ_42_ monomer, and kinetic parameters were derived from 1:1 binding model fits.
*Data are presented as mean ± SEM.
*P < 0.05.

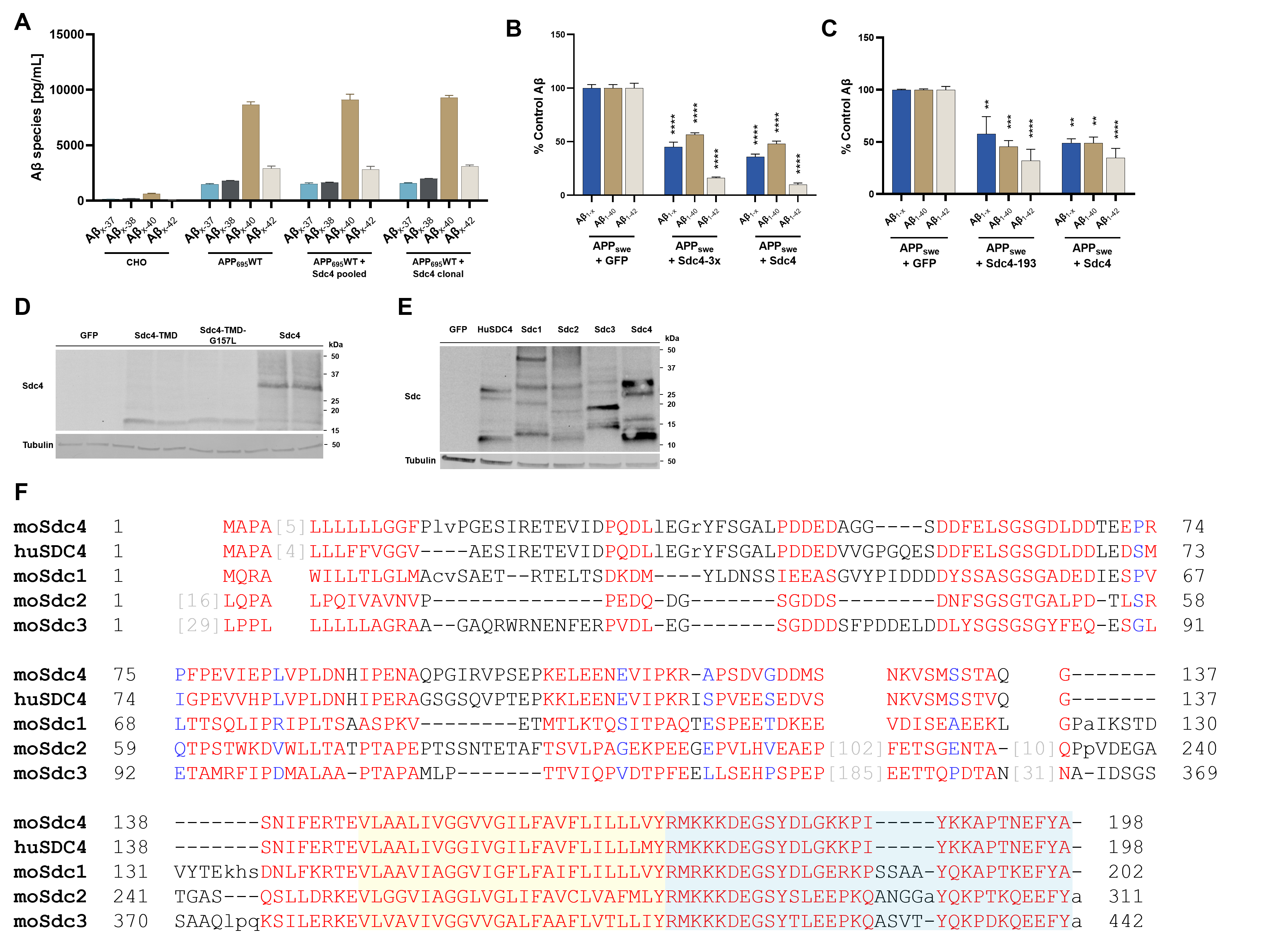

**Supplementary figure 3. Cross-laboratory assay assessment and modulation of Aβ species levels by syndecan-4 variant. (A)** Cross-laboratory assay evaluation using an alternative antibody pair. n = 5 replicates. **(B)** Secreted Aβ species from amyloid precursor protein carrying the K595N and M596L mutations (APP_swe_) co-expressed with Syndecan-4 (Sdc4) or heparan sulfate-deficient Sdc4 mutant harboring S44A, S62A, and S64A substitutions (Sdc4-3x) **(C)** ELISA analysis for detecting Aβ species from the cells co-expressing APP_swe_ with Sdc4 or truncated Sdc4 containing amino acids 1 to 193 (Sdc4-193) **(D)** Expression validation of Sdc4 constructs by immunoblotting. **(E)** Representative immunoblot showing overexpression of Sdc family proteins. **(F)** Multiple sequence alignment of mouse Sdc1-4 and huSDC4. Conserved and variable amino acid residues are shown in alignment format. The transmembrane domain is indicated by the yellow-shaded region, while the cytoplasmic tail is highlighted in cyan. Conserved motifs and sequence features are shown in red, and blue indicates low conservation. Dashes indicate alignment gaps.
*Two-way ANOVA followed by Tukey’s multiple comparisons test
*Data are presented as Mean ± SEM.
**p < 0.01, ***p < 0.001, ****p < 0.0001.

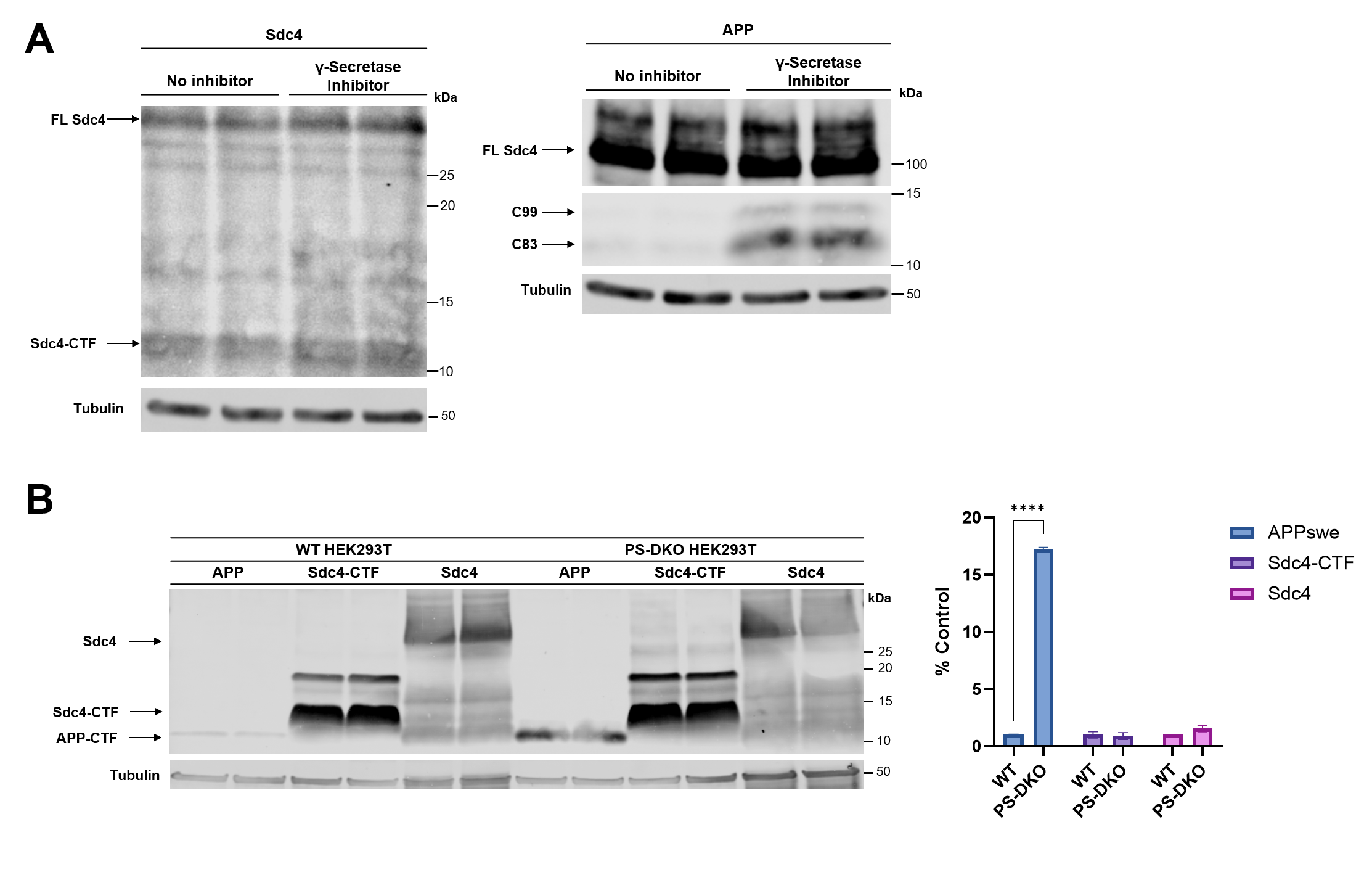
 **Supplementary figure 4. Assessment of γ-secretase–dependent processing of Syndecan-4. (A)** HEK293T cells were transfected with syndecan-4 (Sdc4) and treated with the γ-secretase inhibitor BMS-906024. **(B)** Wild-type (WT) HEK293T and presenilin-1/2 double knockout (PS-DKO) HEK293T cells were transfected with amyloid precursor protein (APP), Sdc4 C-terminal fragment (Sdc4-CTF), or full-length (FL) Sdc4.
*APP-CTF, FL Sdc4, and Sdc4-CTF quantifications normalized to beta-tubulin.
****p < 0.0001.

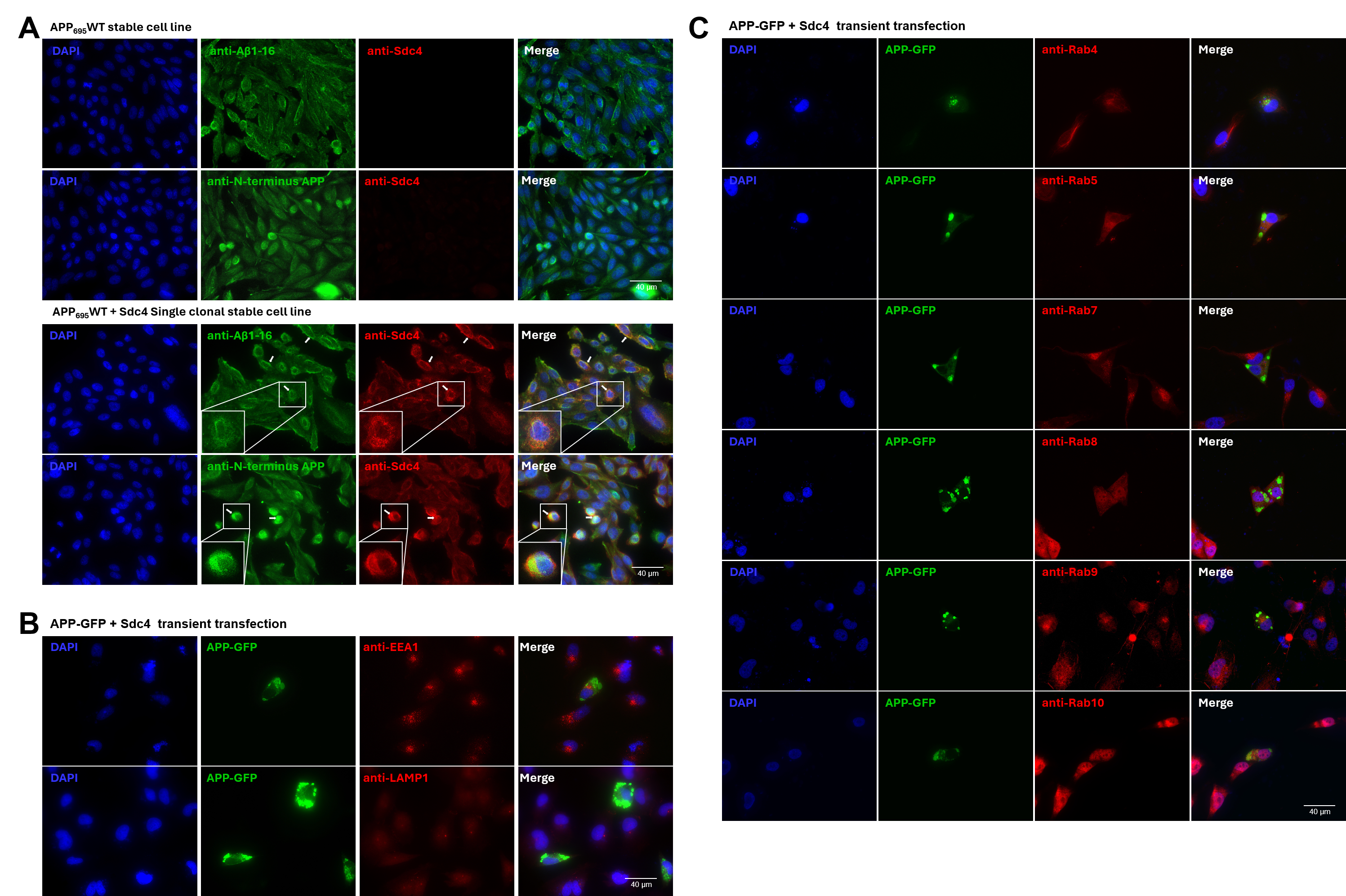

**Supplementary figure 5. Other vesicle markers in H4 cells co-expressing APP and syndecan-4. (A)** Representative immunofluorescence images of CHO stable cell lines expressing APP_695_WT or APP695WT with Sdc4. Cells were immunostained with antibodies against Aβ1–16 (AB5) and the N-terminal region of APP (22C11). **(B)** Distribution of APP-GFP with EEA1 and LAMP1 in H4 cells co-transfected with APP-GFP and syndecan-4. Cells stained for EEA1 and LAMP1, with green representing APP-GFP and red representing the respective markers. **(C)** H4 cells co-transfected with APP-GFP and syndecan-4 were stained for vesicle-associated proteins Rab4, Rab5, Rab7, Rab8, Rab9, and Rab10. APP-GFP is shown in green, and the corresponding markers are displayed in red.
*Nuclei are stained with DAPI (blue).
*Scale bar = 40 μm.

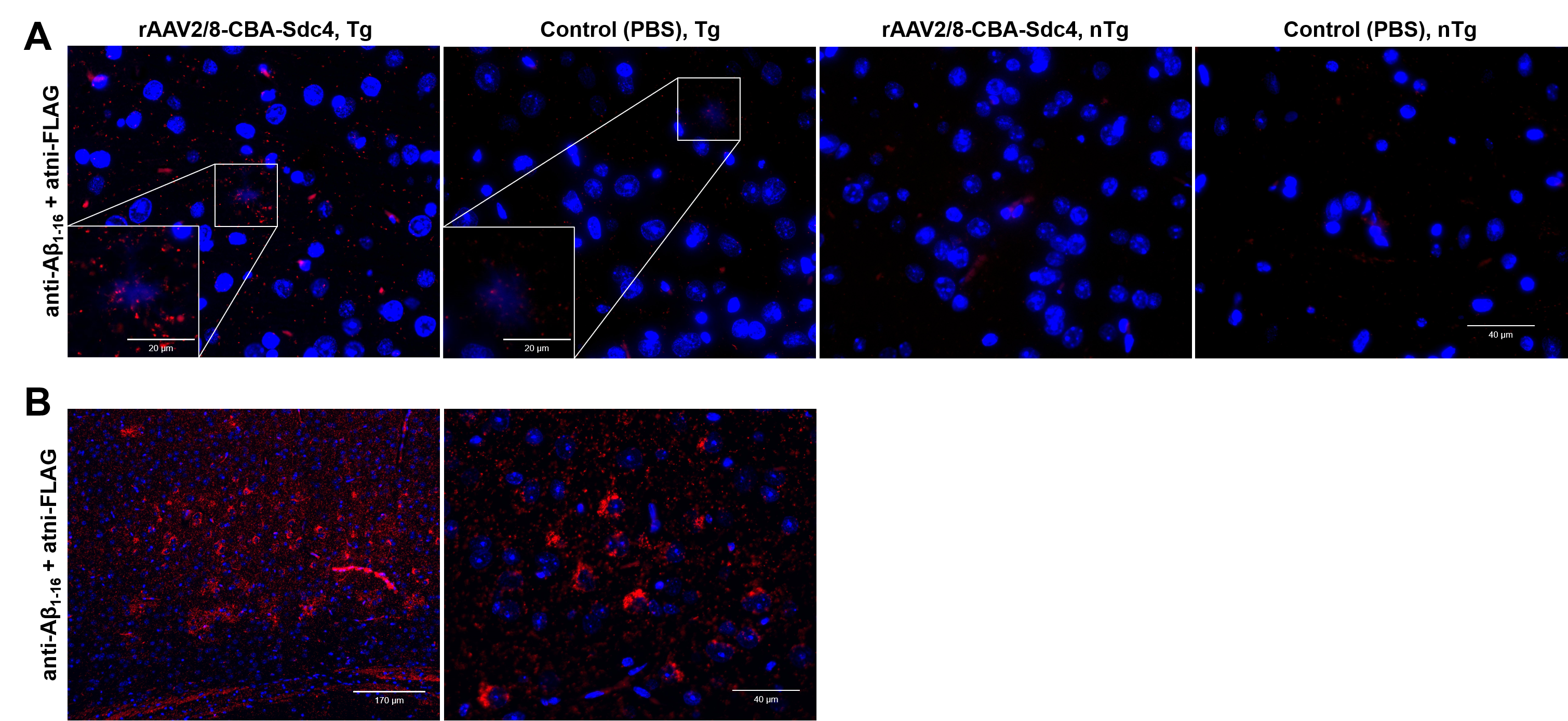

**Supplementary figure 6. PLA detection of syndecan-4 and Aβ in CRND8 mice treated with rAAV2/8-CBA-Sdc4.
(A)** Proximity ligation assay (PLA) was performed in mice brain sections. Red fluorescence indicate PLA signals, reflecting close proximity (<20 nm) between Sdc4 and Aβ. Nuclei were counterstained with DAPI (blue). Scale bar = 40 μm. inset: 20 μm **(B)** Left panel shows a low-magnification overview (scale bar: 170 µm), highlighting the widespread distribution of PLA signals across the tissue. Right panel shows a higher-magnification view (scale bar: 40 µm), illustrating PLA signals concentrated around the nucleus in cells.

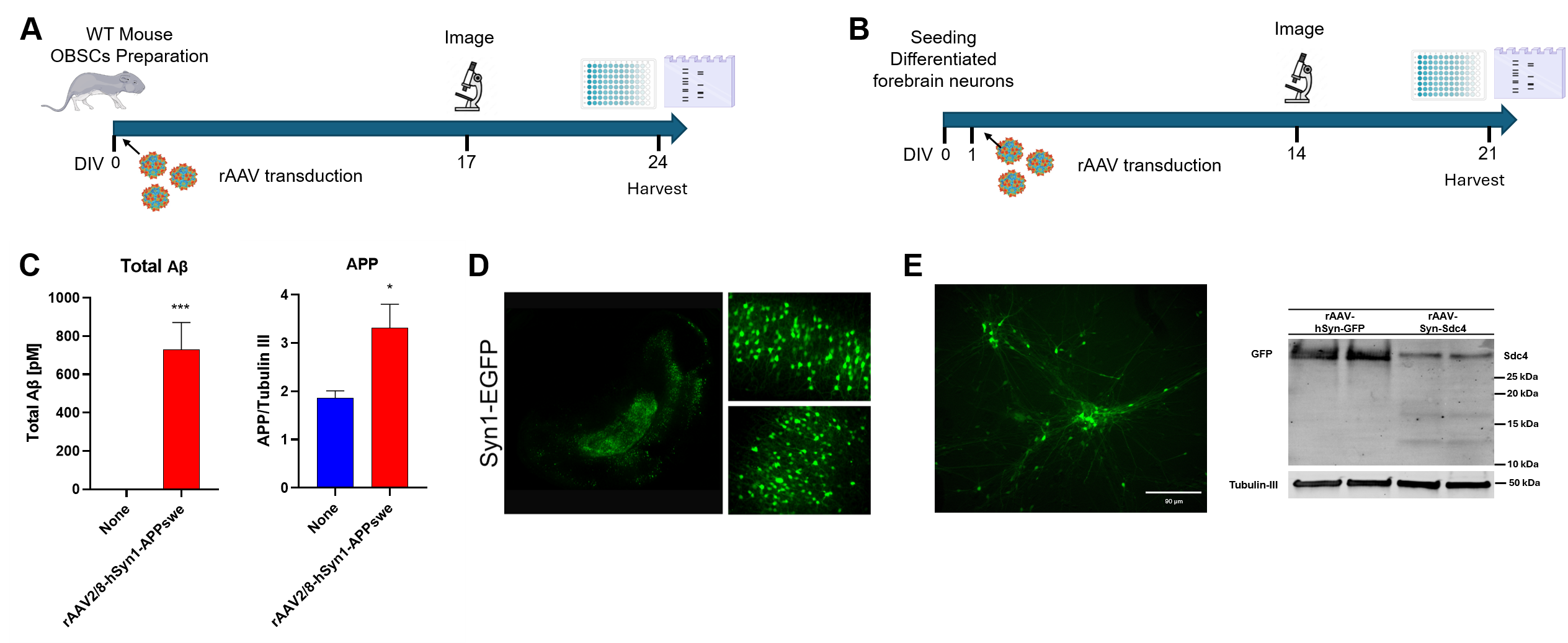
 **Supplementary figure 7. Experimental workflows and biochemical analysis of Aβ levels and APP processing in hiPSC-derived forebrain neurons mouse brain slice cultures using AAV-mediated expression
(A)** Experimental workflow for organotypic mouse brain slice cultures (OBSCs). WT mouse brain slices were prepared and transduced at DIV 0 with rAAV2/8-hSyn1-APP_swe_ and either rAAV-Sdc4 or rAAV-GFP. Depending on the experiment, Sdc4 was expressed under the neuronal Syn1. Slices were imaged at DIV 17, and tissues and conditioned medium were harvested at DIV 24 for biochemical analysis. **(B)** Experimental workflow for human induced pluripotent stem cells-derived neurons (hiPSC-derived neurons). Human iPSC-derived forebrain neurons were co-transduced at DIV 1 with rAAV2/9P31-hSyn1-APP_swe_ and either rAAV2/9P31-hSyn1-GFP or rAAV2/9P31-hSyn1-Sdc4. Neurons were imaged at DIV 14 and harvested at DIV 21 for biochemical analyses. **(C)** Validation of APP_swe_ expression and Aβ production. Transduction with rAAV2/8-hSyn1-APP_swe_ significantly increased total Aβ levels in the conditioned medium (left) and total APP protein levels in lysates (right) compared to non-transduced controls. Statistical significance was determined by Paired comparison test. Data are presented as mean ± SEM. *p < 0.05, ***p < 0.001.n = 8 replicates. **(D)** Representative figures of GFP expressing brain slides. **(E)** Validation of GFP and Sdc4 expression in hiPSC-derived forebrain neurons.
*Scale bar = 90 μm.
*βIII-tubulin served as a loading control.
